# Objective curriculum-guided design of multi-property proteins

**DOI:** 10.64898/2026.05.25.727596

**Authors:** Longying Liu, Jianquan Zhao, Xiaomin Xie, Simeng Xu, Milong Ren, Xinru Zhang, Zaikai He, Fan Liu, Chungong Yu, Kun Wang, Xinglong Wang, Xinmiao Liang, Xianlong Ye, Dongbo Bu, Han Zhou

## Abstract

Designing functional proteins that simultaneously possess multiple biochemical properties remains a significant challenge, as key protein properties, such as solubility, stability, binding affinity, and chemical resistance, are often interdependent or even conflicting. Current approaches typically attempt to jointly optimize multiple functional objectives in one shot, followed by extensive screening to identify rare feasible designs. Here, we introduce OCDesign, an objective curriculum-guided framework for multi-property protein design. OCDesign is based on the objective curriculum principle, which states that the order in which objectives are introduced can shape the accessibility of functional solutions. In each design round of OCDesign, candidate sequences are generated *in silico*, assessed across multiple properties, selected based on Pareto-optimal trade-offs, and experimentally validated, with each experimental stage testing the role of the newly introduced objective within the curriculum. Using antibody-binding protein A as a model system, we show that one-shot optimization fails to yield functional designs, whereas a staged curriculum—progressing from solubility and structural consistency to binding affinity, and then to alkaline resistance—enables the design of proteins possessing multiple desired properties through substantially fewer wet-lab experiments. These results establish OCDesign as a practical computational–experimental strategy for organizing and integrating multiple objectives in protein design, and suggest that objective ordering is a key determinant of accessibility in high-dimensional design spaces.

## 1 Introduction

Designing proteins with desired functions is a central goal of molecular biology and biotechnology [1–7]. Recent advances in deep learning have enabled accurate prediction of protein structures and the generation of sequences that fold into predefined conformations [8]; however, designing proteins that simultaneously possess multiple functional properties remains a major challenge.

The challenge is exemplified by the design of antibody-binding proteins, especially *Staphylococcal* protein A [9]. Protein A consists of five highly homologous domains connected by short linkers (denoted as domains A, B, C, D, and E; Fig. 1**a**). These domains can specifically bind to the Fc region of certain antibodies (e.g., IgG; Fig. S1), enabling the capture and purification of antibodies using protein A as an affinity ligand or a filler in chromatography columns [10–13]. In these cases, an idealized protein A is expected to simultaneously possess multiple biochemical properties, including high stability, solubility, appropriate binding affinity, and strong alkaline resistance [13–17]. These properties are interdependent or even conflicting; consequently, the optimization of one property might degrade others, and the simultaneous optimization of multiple properties requires intrinsic trade-offs.

**Fig. 1:**
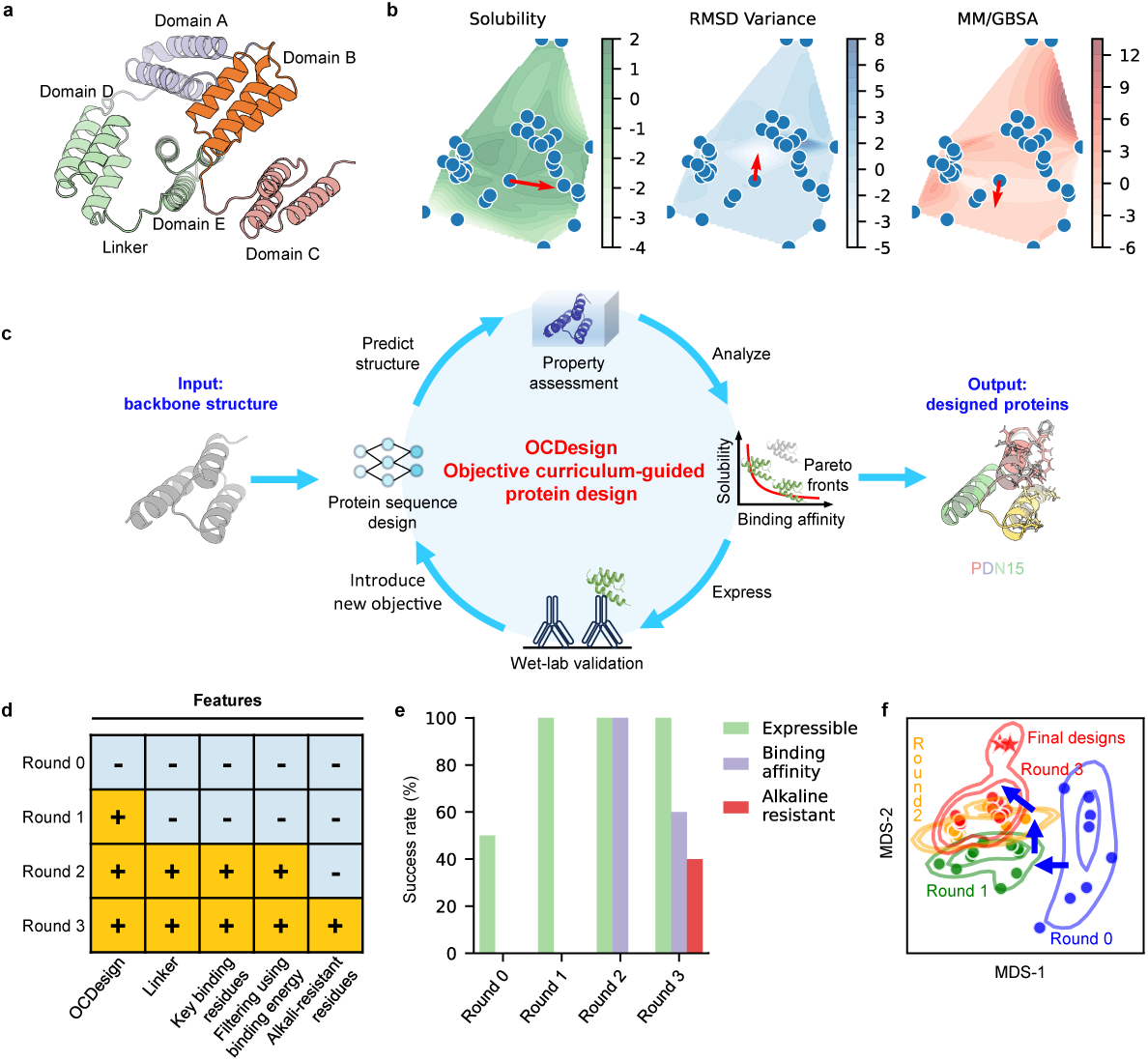
Objective curriculum-guided framework for multi-property protein design. **a,** Protein A consists of five highly homologous domains that are connected by short linkers. **b,** Design landscapes colored by estimated solubility, stability, and binding affinity (measured using MM/GBSA). Property gradients (shown as arrows) vary across objectives, as exemplified by PDN4. The landscapes were generated by multidimensional scaling based on sequence similarity among designs. **c,** Overview of the objective curriculum-guided design framework OCDesign. OCDesign integrates sequence design, *in silico* assessment of biochemical properties, Pareto-optimal selection, and wet-lab experimental validation to test the role of objectives within the curriculum. OCDesign introduced objectives in a predefined order and finally obtained two designs with the desired properties. **d, e,** The features considered in each design round and the corresponding success rates. **f,** OCDesign reshapes the design space and navigates the search dynamics (shown as trajectory arrows in blue), finally enabling access to the successful designs with intended properties (labeled as stars)

The design and optimization of functional proteins can be accomplished by directed evolution, which mimics natural selection in labs to engineer proteins with desired properties through iterative rounds of mutation and screening [18, 19]. This strategy has achieved great success but always incurs high time and resource costs, requires specialized expertise, and is limited by local exploration of protein sequence space—the final designs generally mutate only a few residues, leading to high sequence identity over 0.90 [20]. In parallel, computational approaches have made significant advances in designing protein structures and sequences by learning sequence-structure relationships [8, 21–26]. However, these approaches primarily optimize structural fidelity and often fail to achieve robust functional performance.

A straightforward strategy for multi-property protein design is one-shot optimization, which simultaneously optimizes multiple objectives, followed by extensive screening to identify feasible designs. However, this strategy often fails or requires many trials [27], suggesting that multi-property protein design is determined not only by which objectives are considered but also by how they are introduced into the design process. More broadly, successful protein modeling approaches have often relied on staged optimization strategies, in which constraints are introduced or emphasized progressively during the search for native-like structures [28, 29], implying that the ordering of constraints may also play an important role in navigating complex design landscapes.

Here, we introduce OCDesign, an objective curriculum-guided framework for multi-property protein design. OCDesign is guided by the objective curriculum principle (OCP), which posits that objective ordering plays a central role in shaping the search dynamics of multi-property design. Instead of optimizing all objectives in one shot, OCDesign organizes objectives into a curriculum in which early-stage objectives establish basic properties (i.e., residue compatibility and protein foldability), whereas later-stage objectives introduce increasingly specific functional requirements. This staged organization progressively constrains and refines the feasible design space, facilitating the integration of multiple properties.

We implemented OCDesign by integrating computational sequence generation, *in silico* property assessment, Pareto-optimal selection, and wet-lab validation. Using antibody-binding protein A as a model system, we systematically compared one-shot optimization with a staged objective curriculum. One-shot optimization failed to produce functional proteins, whereas the staged curriculum—progressing from solubility and structural consistency to binding affinity, and then to alkaline resistance—enabled successful multi-property design with substantially fewer wet-lab experiments.

Notably, the final design strategy that emerges from this process is conceptually simple, relying on fixing key residues and optimizing sequences under multiple objectives. However, this simplicity is not apparent *a priori*, emerging only after navigating the complex and highly constrained design landscape through iterative exploration. Rather than proposing a complex method, our work identifies a simple, effective strategy and explains why it works only when objectives are ordered in a specific way.

Together, these results establish OCDesign as a practical strategy for converting multi-property protein design from one-shot joint optimization into staged, hypothesis-driven objective integration.

## 2 Results

### 2.1 One-shot design strategy fails to satisfy multiple objectives simultaneously

We first asked whether multiple properties could be achieved through one-shot optimization. To this end, we conducted a preliminary design campaign that attempts to optimize three properties (i.e., solubility, structural self-consistency, and binding affinity) in a single stage. The details are provided in the Supplementary material. To examine the effects of objectives, we also repeated the entire design process, optimizing only two objectives: solubility and structural self-consistency.

We finally acquired 19 designs and selected six with high values of the combined objective function for wet-lab validation, including four obtained by optimizing three properties and two by optimizing only two properties. The sequences and properties of these designs are listed as “Round 0” in Tables S1, S2, and S3. However, experimental characterization of the six designs revealed that none binds to the target antibody, and three are insoluble or poorly expressed. This finding is consistent with recent studies on binder design, which revealed the challenge of achieving multiple properties in a single stage, leading to a low success rate and often requiring many trials [27].

These results highlight a fundamental limitation of one-shot multi-property optimization: although computational models can identify sequences that satisfy each objective individually, they fail to capture the complex dependencies required for functional integration, thereby ignoring the path-dependent nature of navigating the feasible design space. We present solid evidence of the conflicts among multiple properties in Fig. 1**b**: different properties typically exhibit opposite directions of improvement; thus, improving one property often degrades others.

Together, these findings suggest that the feasible design space, defined by the intersection of multiple constraints, is highly restricted and difficult to access directly, thereby motivating the need for an objective curriculum.

### 2.2 Residue-compatibility constraints establish a feasible but non-functional design space

Under the objective curriculum, we began by designing proteins with only two properties (i.e., solubility and structural self-consistency) rather than attempting to achieve multiple properties in a single stage. Previous studies have shown that the residues of a protein should possess considerable inter-residue compatibilities, which are critical to its solubility and stability [30, 31]. Hence, we hypothesized that imposing residue-compatibility constraints will facilitate the establishment of a feasible design space for foldability and expression.

To test this hypothesis, we designed protein sequences by enforcing interresidue compatibility using two deep learning models, i.e., NeuralMRF and ProDESIGN-LE [23]. In particular, NeuralMRF extracts the inter-residue dependency from the native 3D structure of the natural protein A using a message-passing neural network, and then decodes the dependency into protein sequences using the Markov random field (MRF) technique [32, 33]. The details of NeuralMRF are provided in the Supplementary material. To enhance design diversity, we also applied ProDESIGN-LE, a protein design approach that leverages residue preferences for amino acid types derived from the local structural environment around each residue [23]. It should be noted that both sequence-generating approaches can begin by fixing certain key residues, thereby enabling the enhancement of additional properties in subsequent design stages.

We used these two approaches to design sequences based on the B-domain structure of protein A, and then constructed full-length proteins that directly cascade four copies of the designed domain sequence, with no linkers between them. This way, we acquired a total of 10 full-length designed proteins, including five from NeuralMRF and five from ProDESIGN-LE. Next, we assessed solubility and structural self-consistency of these designs *in silico*. The sequence and properties of these designs are provided as “Round 1” in Table S4 and S5.

To reduce the need for wet-lab experiments, we computed and focused on Pareto fronts of the initial designs, which offer the best trade-offs between the two properties. As shown in Fig. 2, PDN3, PDN4, and PDN5 are Pareto fronts that dominate the other designs by NeuralMRF, and PDN6, PDN8, and PDN10 are Pareto fronts that dominate the other designs by ProDESIGN-LE. Here, we selected one design by NeuralMRF (PDN4, solubility: 0.920, TM-score: 0.963) and one design by ProDESIGN-LE (PDN8, solubility: 0.926, TM-score: 0.968) for further wet-lab validation.

**Fig. 2:**
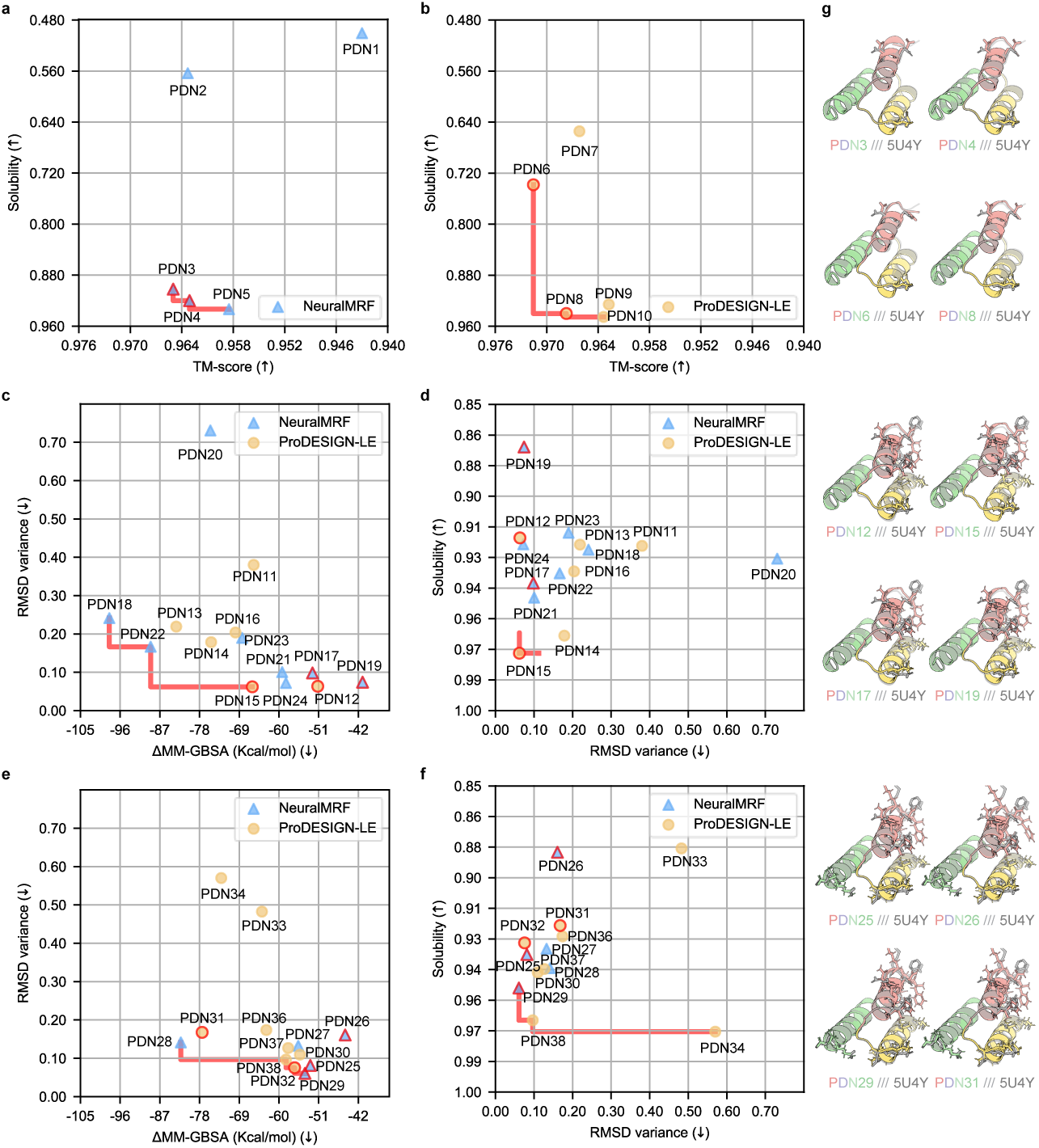
Computing the Pareto front designs that offer the best trade-offs across multiple properties. **a, b,** The designs obtained from round 1 using NeuralMRF (**a**) and ProDESIGN-LE (**b**). Hereinafter, the Pareto fronts are connected and thus labeled by a line, and the designs selected for further wet-lab validation are outlined. **c, d,** Two 2D views of the round 2 designs. OCDesign assesses three properties of the designs. For improved clarity, we visualize these designs in two 2D views rather than a single 3D view. **e, f,** Two 2D views of the round 3 designs. **g,** The predicted structures of the selected designs for wet-lab validation.

The wet-lab experimental results revealed that the selected designs are all soluble and successfully expressed (Table S6 and Fig. 3**a**), supporting the hypothesis that residue-compatibility constraints are helpful for achieving basic physicochemical properties. Furthermore, we selected a design (PDN8) as a representative for secondary-structure characterization using circular dichroism (CD) spectroscopy, which indicates that its secondary structure is in good agreement with the intended structure (Fig. 4**a**).

**Fig. 3:**
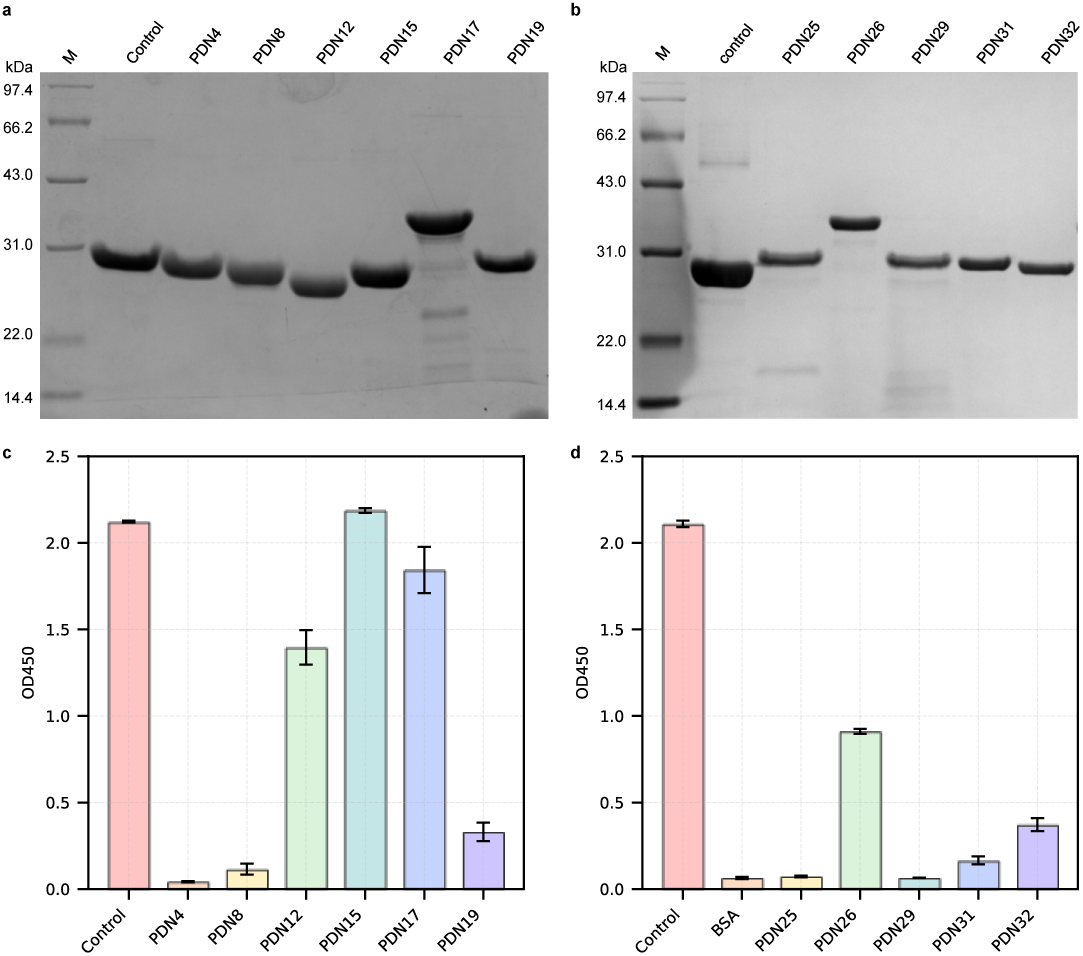
Solubility and binding affinity of the designs by OCDesign. **a, b,** Characterizing solubility of the designs using SDS-PAGE. **c, d,** Assessing IgG-binding affinity of the designs using ELISA.

**Fig. 4:**
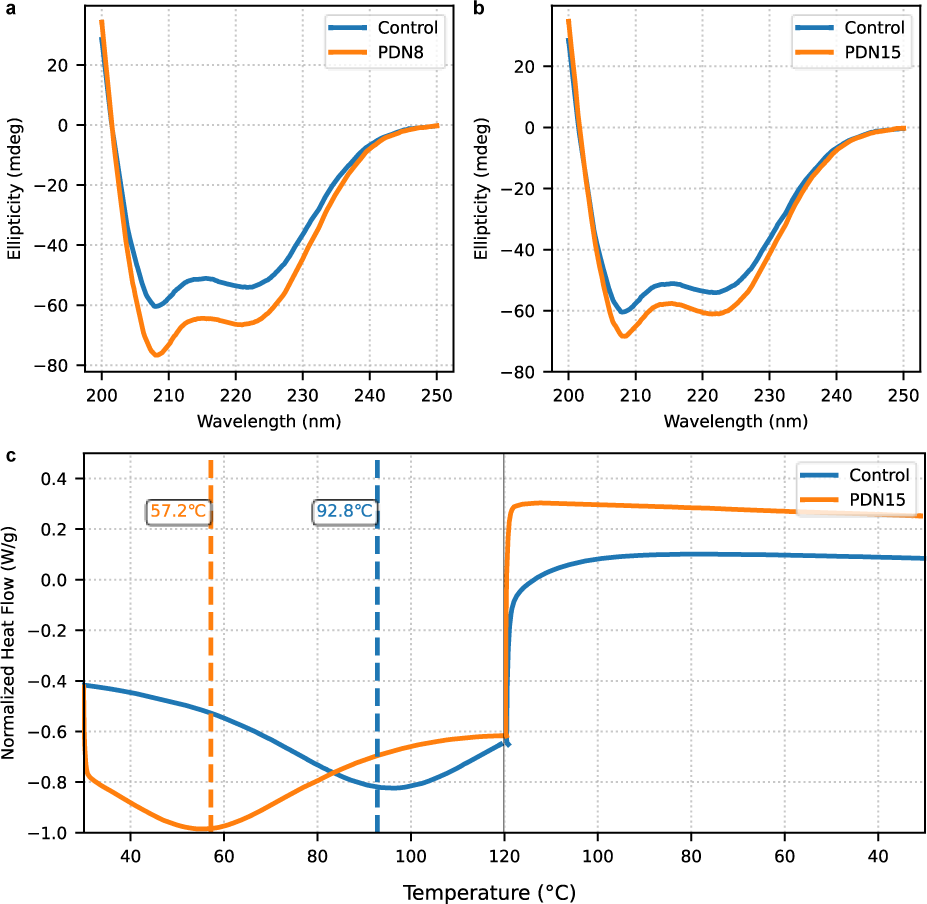
Structural characterization of two designs, PDN8 and PDN15, using circular dichroism. Here, we use two Pareto fronts (PDN8 and PDN15) as representatives. **a, b,** Circular dichroism spectra of the designs PDN8 and PDN15 (orange) from 200 to 250 nm at 25 °C compared to the control protein (blue). Both exhibit characteristic “W”-shaped spectra with minima at 208 nm and 222 nm. **c,** Thermal stability analysis of the two proteins using differential scanning calorimetry. The designed protein PDN15 shows a denaturation temperature of 57.2 °C, whereas the natural protein A shows a denaturation temperature of 92.8 °C.

In addition, these results also confirmed the effectiveness of both NeuralMRF and ProDESIGN-LE in generating soluble proteins. Consequently, to increase design diversity, we applied both approaches in subsequent design rounds and processed their designed sequences using the same pipeline. However, these designs failed to exhibit antibody-binding activity (Fig. 3**c**), indicating that residue-compatibility constraints alone are insufficient for functional performance.

Together, these findings are consistent with the first stage of the objective curriculum principle, in which early objectives constrain the design space toward foldable and expressible proteins, without guaranteeing their functional performance.

### 2.3 Interface-specific constraints enable the emergence of binding

We then hypothesized that antibody binding requires the introduction of interface-specific constraints that enforce precise residue-level interactions. We also hypothesized that the failure in round 1 comes from a lack of linkers to connect the designed domains into a full-length protein.

To test the first hypothesis, we identified key residues involved in antibody binding and incorporated them into the design process. Specifically, we first identified such key residues by molecular dynamics (MD) simulation analysis of the protein A-antibody complex (PDB ID: 5U4Y) using the Schrödinger suite [34–37]. Seven residues of the wild-type protein A were found to tightly interact with the antibody, including Phe1, Gln6, Asn7, Phe9, Tyr10, Glu20, and Asn24 (Fig. S1). Specifically, residues Phe1, Phe9, and Tyr10 of protein A form *π − π* stacking interactions with residues Tyr200, His74, and His197/His199 of the antibody, respectively. In addition, Gln6 forms three hydrogen bonds with Leu15, Ile17, and Asn198 of the antibody. Asn7 and Asn24 also form hydrogen bonds with Asn198 and Gln75 of the antibody, respectively. Glu20 interacts with Lys81 on the antibody via a hydrogen bond and a salt bridge simultaneously. Mechanistically, the binding energy of the complex is primarily localized to these critical interaction hotspots. These findings are also consistent with the previous study on sequence conservation [38].

Next, we ran NeuralMRF and ProDESIGN-LE with these key residues fixed, yielding a total of 14 designs (Round 2, Table S4). To examine whether these key residues can facilitate binding, we performed MD simulations on the complex formed by a designed protein and the antibody. The MD simulations report structural variations, and the binding energies calculated via the MM/GBSA method provide approximate measures of the binding strength (Fig. 5, S2, S3, S4, S5 and Table S7).

**Fig. 5:**
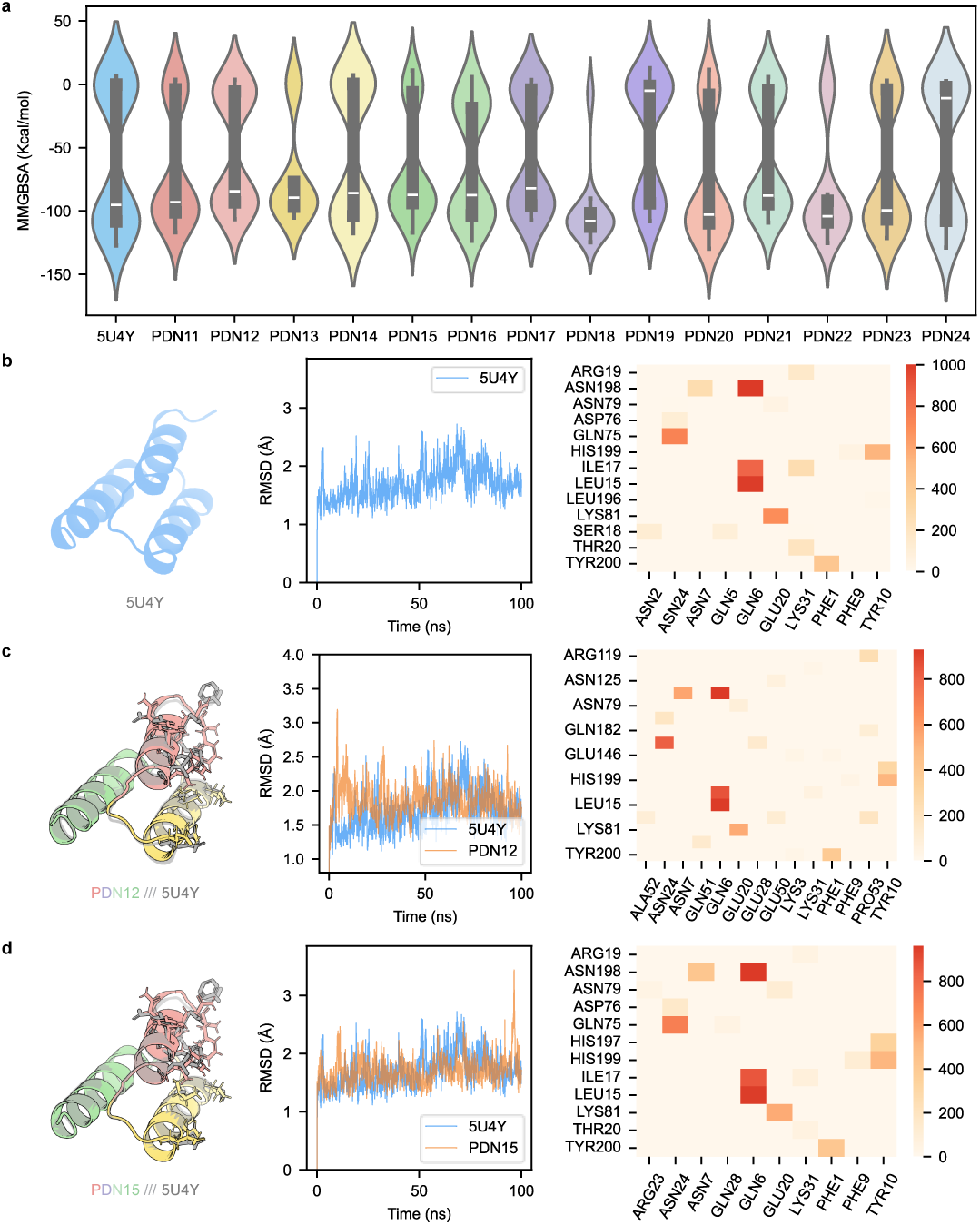
Deriving the interface-specific constraints from molecular dynamics simulation and imposing these constraints in protein design. We first performed MD simulations (100 ns) by running Schrödinger on the protein A-IgG complex. From the MD trajectory, we identified key binding residues as interface-specific constraints and imposed these constraints in subsequent protein design. We present two designs (PDN12 and PDN15) here and provide the other designs in the Supplementary material. **a,** The distribution of binding energy (MM/GBSA) of complexes calculated using the final 10 ns trajectory. **b, c, d,** The MD simulation trajectories and key binding residues of the complexes formed using the natural protein A, PDN12, and PDN15. The binding residues are shown using a matrix with columns denoting IgG residues and rows denoting residues of the natural protein A or designs.

Then, we computed Pareto fronts from the initial 14 designs that best trade off three properties: solubility, structural stability during MD simulation (measured by RMSD variance), and binding energy (measured by ΔMM/GBSA). Here, we used structural variation rather than structural self-consistency, as it carries more information, especially molecular dynamics. As shown in Table S7 and Fig. 2**c-d**, the 14 initial designs form four Pareto fronts: PDN14, PDN15, PDN18, and PDN22. We selected one Pareto front (PDN15) for wet-lab validation. To examine the effectiveness of Pareto-front selection, we also selected three non-Pareto-front designs (PDN12, PDN17, and PDN19), which have structural stability comparable to PDN15 and high binding energy.

To test the second hypothesis, we performed molecular dynamics analysis of the complex formed by the target antibody and the full-length proteins and investigated the effects of linkers. Our molecular dynamics analysis suggests that the full-length proteins, which directly cascade four copies of the designed domain sequence with no linkers inside, exhibit substantial rigidity. This rigidity reduces the number of antibody-bound domains that can avoid steric clashes with the plate, thereby decreasing the likelihood of effective binding (Fig. S6). In contrast, the full-length proteins, when inserted with linkers between adjacent domains, exhibit relatively high flexibility, thereby reducing the likelihood of clashing and increasing the likelihood of binding. Consequently, in this round, we use full-length proteins with inserted linkers. Here, each linker is a 5-mer peptide, “KVDAK”, derived from the natural protein A.

Wet-lab experimental characterization of the resulting four full-length proteins revealed that three of them, i.e., PDN12, PDN15, and PDN17, exhibit considerable binding affinity with the target antibody (Fig. 3 and Table S8). Notably, PDN15 shows the highest IgG-binding affinity (OD_450_: 2.15), supporting the effectiveness of Pareto fronts over non-Pareto front designs. Further analysis of PDN15 using circular dichroism spectroscopy reveals that its secondary structure closely agrees with the natural protein A (Fig. 4**b**).

These results support the hypothesis that binding emerges only after introducing constraints that govern specific intermolecular interactions, and binding can be boosted after introducing constraints that facilitate these interactions. Importantly, the emergence of binding is consistent with the objective curriculum principle, which posits that objectives requiring specific intermolecular interactions should be introduced after basic compatibility and foldability have been established. Both interface-specific constraints and linker-introduced structural flexibility likely contribute to binding, although their individual contributions may be intertwined.

Nevertheless, these designs remain susceptible to alkaline degradation (Fig. 6), indicating that additional constraints are required to achieve full functional robustness.

**Fig. 6:**
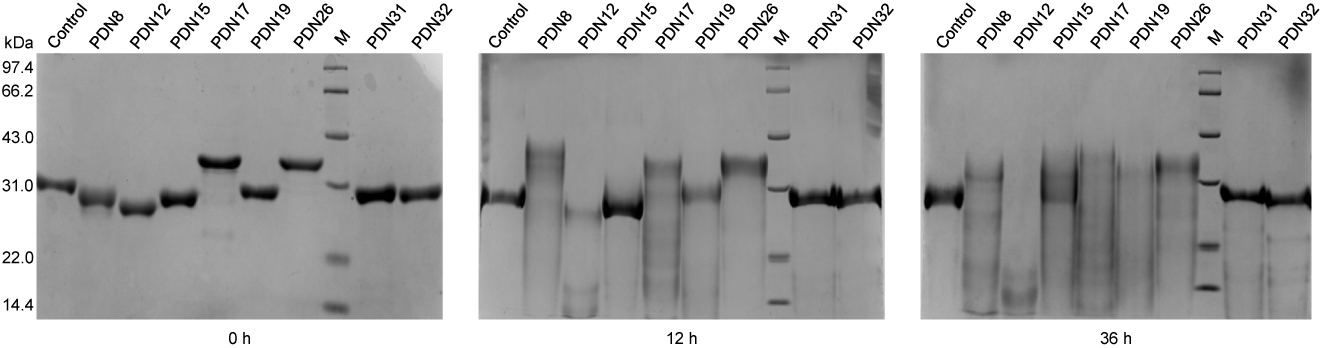
Characterizing alkaline resistance of the designed proteins using SDS-PAGE. To investigate the alkaline resistance of the designed proteins, we continuously treated them with 0.5 mol/L NaOH solution (after 0, 12, and 36 hours) and examined their integrity using SDS-PAGE. The data acquired after 24- and 48-hour treatments are provided in the Supplementary material.

### 2.4 Environmental robustness requires additional orthogonal constraints

We next asked whether environmental robustness, such as alkaline resistance, can be achieved by extending the objective curriculum further. We hypothesized that such properties are governed by constraint classes that are partially independent of those controlling structure, solubility, and binding.

To test this, we introduced mutations associated with alkaline resistance while preserving previously established constraints. Previous studies have shown that residue mutations N2A, N19T, G25W, and Q36V contribute greatly to alkaline resistance [14, 39]. Here, we fixed these residues along with the key binding residues, and re-executed OCDesign, yielding an additional 14 designs, including 7 by NeuralMRF and 7 by ProDESIGN-LE (Round 3, Table S4).

Analysis of these 14 initial designs reveals five Pareto fronts, i.e., PDN28, PDN29, PDN32, PDN34, and PDN38 (Table S9, Fig. 2**e, f**). We selected two Pareto fronts (PDN29 and PDN32) and three non-Pareto fronts (PDN25, PDN26, and PDN31) for wet-lab validation.

Wet-lab experimental results reveal that all of the five designs are soluble, and three of them (PDN26, PDN31, and PDN32) show considerable IgG-binding affinity (Fig. 3 and Table S10). Furthermore, we evaluated the alkaline resistance of these three proteins, i.e., whether they retain considerable binding affinity under alkaline conditions. As shown in Fig. 6, two of them, PDN31 and PDN32, exhibit significant alkaline resistance, indicating that structural integrity is maintained under alkaline conditions while retaining binding affinity. In addition, the Pareto optimization remained useful: one out of the two Pareto fronts (i.e., PDN32) exhibits considerable alkaline resistance, whereas only one out of the three non-Pareto fronts (i.e., PDN31) shows similar functional performance.

Together, these findings support the hypothesis that environmental robustness requires additional orthogonal constraints, and that such constraints can be integrated only after structural and functional requirements have been satisfied. This hierarchical organization of the constraints enables sequential layering of the objectives operating at different scales.

### 2.5 Objective curriculum progressively reshapes the design landscape

Finally, we examined how the sequential introduction of objectives influences the overall design process.

To investigate the design space and search dynamics, we depicted the designs obtained across all rounds in Fig. 1**f**. The designs acquired in round 0 are relatively dispersed, indicating randomness in the sequence search and a low success rate. In contrast, when the objective curriculum is applied, the resulting designs are significantly more compact, suggesting a narrower design space. In addition, the design space systematically shifts as more objectives are introduced, ultimately enabling the identification of successful designs. These results reflect a gradual reshaping of the protein design landscape, in which experimentally validated constraints reduce the accessible sequence space.

The reshaping of the design space results in a considerable increase in the success rate of designs that satisfy multiple properties across successive stages. As shown in Fig. 1**e**, the two designs selected in round 1 (PDN4 and PDN8) are both expressible. In round 2, the four selected designs (PDN12, PDN15, PDN17, and PDN19) are all expressible and exhibit considerable IgG-binding affinity. In round 3, three of the five selected designs (PDN25, PDN26, PDN29, PDN31, and PDN32) exhibit IgG-binding affinity, and two of them (PDN31 and PDN32) also show strong alkaline resistance, achieving a 40% success rate in designing proteins that possess all three properties simultaneously. These findings support the effectiveness of an objective curriculum in guiding functional protein design while reducing the number of wet-lab validation experiments.

Furthermore, Fig. 1**e** also reveals a systematic shift in the distribution of protein properties across successive design rounds. Early designs satisfy only basic physicochemical constraints, whereas later designs achieve simultaneous optimization of solubility, binding affinity, and alkaline resistance. The observed progression—from foldability to binding to environmental robustness—suggests that each stage progressively refines the feasible solution space towards the final successful designs. Importantly, the designs selected based on Pareto-optimal trade-offs consistently outperformed non-Pareto candidates, highlighting the effectiveness of multi-objective selection in constrained design spaces.

We expressed the designed proteins in *Escherichia coli* and examined whether they possess the desired properties. As shown in Fig. 3, 6, S7, and Table S11, the final designs acquired in round 3, PDN31 and PDN32, have high solubility and alkaline resistance, as well as appropriate IgG-binding affinity, making them promising substitutes for the natural protein A as fillers for antibody purification. The functional characterization of the other designs is provided in the Supplementary material.

Notably, the final design strategy is conceptually simple, relying on fixing key residues and optimizing sequences under multiple objectives. However, the counterpart one-shot optimization of multiple objectives failed, indicating that the effectiveness of this approach critically depends on the objective curriculum. These findings provide evidence that objective ordering plays a central role in shaping the search dynamics of multi-property protein design.

Together, these results demonstrate that iterative incorporation of experimentally validated constraints enables efficient navigation of complex multi-property landscapes and leads to the discovery of functional proteins that cannot be obtained through one-shot or purely computational approaches.

## 3 Discussion

This study presents OCDesign, an objective curriculum-guided framework for multi-property protein design. By integrating computational sequence generation, Pareto-based multi-objective optimization, and wet-lab validation, OCDesign provides a practical strategy for organizing multiple design objectives that are difficult to optimize simultaneously. Our results show that successful multi-property design depends not only on the objectives to be optimized but also on the order in which they are introduced. This organization improves design efficiency while allowing each experimental stage to test how newly introduced objectives interact with previously established constraints.

Essentially, the feasible design space under multiple constraints is not only small but also structured, such that certain regions are not merely difficult to access but conditionally inaccessible unless preceding constraints are satisfied. Unlike traditional staged optimization, which is often heuristic and method-specific, this work identifies objective ordering itself as an independent axis governing accessibility in high-dimensional optimization. In other words, the feasibility under multiple objectives is path-dependent, rather than solely determined by the objectives themselves.

The results also reveal the effectiveness of Pareto-optimal selection for trading off multiple objectives. However, the Pareto selection alone is insufficient without appropriate objective ordering, as evidenced by the failure of the one-shot optimization strategy.

A notable feature of our results is that late-stage objectives can be incorporated without disrupting the properties established in earlier stages. This feature stems from the hierarchical nature of constraints imposed by our objective curriculum. In particular, early-stage objectives, such as solubility and structural consistency, impose coarse-grained constraints supporting foldability and expression. In contrast, later-stage objectives, such as binding affinity and alkaline resistance, introduce specific interaction requirements. Under this order, late-stage constraints refine rather than overwrite earlier solutions, therefore enabling sequential integration of multiple properties.

Interestingly, similar ideas have implicitly appeared in protein structure prediction: classical approaches search for the native structure using a staged optimization strategy, which emphasizes local structure formation (e.g., secondary structure elements) in early stages and global topology (e.g., interactions among these elements) in later stages [28]. While such approaches were not typically framed in terms of an explicit curriculum over structural objectives, their success suggests that the ordering of structural features reflects the essence of protein folding and thus can influence the accessibility to native structures. In this sense, our objective curriculum principle can be viewed as a generalization of these staged optimization strategies to multi-property protein design.

An important implication of our design framework is that wet-lab experiments serve not only to validate the acquired designs but also to test how newly-introduced objectives interact with the constraints established in previous stages. Notably, in the present study, the objective curriculum is predefined based on experts’ domain knowledge, providing a controlled setting to isolate the causal role of objective ordering.

The present study still has several limitations. First, the objective curriculum used in this study is manually specified and may not be optimal. An idealized objective ordering should reflect intrinsic constraint structure rather than heuristic choices. Hence, developing approaches to find the optimal curriculum or dynamically construct it from wet-lab validation results remains an important direction for future work. Second, computational models used to assess protein properties are imperfect and may bias design selection. Third, our experiments focus on a single protein system and protein sequence design; broader validation across diverse proteins and protein sequence-structure co-design will facilitate the assessment of the method’s generality.

Despite these limitations, OCDesign provides a generalizable strategy for converting multi-property protein design from one-shot joint optimization into staged, hypothesis-driven objective integration. More broadly, our results suggest that accessibility in high-dimensional design spaces is governed not only by whether objectives are compatible, but also by the trajectory through which constraints are imposed.

## 4 Methods

The OCDesign framework designs multi-property proteins using staged optimization under a pre-defined objective curriculum. In the first round, OCDesign aims to design proteins with only two properties: solubility and structural self-consistency. In the second round, an additional property (binding affinity) was introduced into the optimization process. In the third round, OCDesign considers all four properties, including solubility, stability, binding affinity, and alkaline resistance.

Each design round consists of four steps, which are depicted in Fig. 1**c** and described below:

*(i) Protein generation:* The residues of a protein should possess considerable inter-residue compatibilities, which are critical for the protein’s solubility and stability [30, 31]. To generate such protein sequences, we developed a deep learning model (NeuralMRF) that explicitly learns and exploits the compatibilities (See Supplementary Methods). To test the generality of our framework and boost design diversity, we also applied ProDESIGN-LE to generate sequences [23]. Both sequence-generating approaches can begin by fixing several key residues, thereby enabling sequential introduction of objectives in the design process.
*(ii) Multi-objective optimization:* In round 1, we *in silico* assessed two biochemical properties of the designed proteins, including protein solubility predicted using SoluProt [40], and structural self-consistency, i.e., the consistency between their predicted structures and the native structure of natural protein A. In rounds 2 and 3, we further assessed the protein-antibody complexes, each consisting of a designed protein and the target antibody. Specifically, we evaluated stability, binding affinity, and key residue interactions of the complexes by running Schrödinger suite [34–37] on the predicted complex structures (see Supplementary Methods for details). To identify the designs that simultaneously optimize multiple biochemical properties, we computed the Pareto fronts, i.e., the designs that make the best trade-off among these properties, as these designs cannot be improved in one property without degrading any other properties [41]. These Pareto front designs were selected for subsequent wet-lab validation.
*(iii) Wet-lab validation:* We performed wet-lab experiments to validate these selected designs, i.e., expressing them *in vitro*, quantitatively analyzing their binding affinity to the target antibody, and assessing their alkaline resistance. More importantly, the wet-lab experiments also test the role of objectives within the curriculum. Each stage of experimentation evaluates a stage-specific hypothesis, providing evidence on how newly-introduced objectives interact with the constraints established in previous stages. We analyzed wet-lab experimental results to gain experience in sequence generation and selection, and exploited these validation results to improve future designs.
*(iv) Introducing new objective according to curriculum:* In each round, OCDesign introduces a new objective into the optimization process and starts a new design round. The objective ordering was chosen to progressively introduce more specific functional constraints, i.e., from solubility and structural consistency to binding affinity and, finally, to alkaline resistance. After three rounds, OCDesign produces the final designs expected to possess the intended properties.

The details of *in silico* assessment of protein properties and wet-lab experiments are provided in the Supplementary material.

## Availability

The source code and data for OCDesign are freely available at https://github.com/bigict/OCDesign.

## Acknowledgments

We acknowledge the support from the National Key Research and Development Program of China (2024YFC3405501), the National Natural Science Foundation of China (32271297), the Jiangxi Provincial Key Research and Development Program (2025BAC250061), and the Special Fund for Digital Economy on Budgetary Infrastructure Investment of Jiangxi Province (2506-360000-04-04-511042). The Computer-X center at the Institute of Computing Technology supported the numerical calculations in this study. We also thank Dr. Haicang Zhang for helpful discussions.

## Supplementary material

### 1 Results

#### 1.1 Functional characterization of the designed proteins

We expressed the designed proteins in *Escherichia coli* and examined whether they possess the desired properties as follows.

##### (i) Protein solubility

We assessed the solubility of the designed proteins using SDS-PAGE. As shown in Fig. 3**a** and **b**, the final designs (PDN25, PDN26, PDN29, PDN31, and PDN32), together with the Pareto fronts selected in round 1 (PDN4 and PDN8) and round 2 (PDN12, PDN15, PDN17, and PDN19), exhibit high expression comparable with the natural protein A. Indepth examination suggested that these designs were recovered exclusively in the soluble fraction (Fig. S8, S9, and S10). We further conducted SECHPLC analysis, which revealed a high purity of the refined proteins (99.37%, Fig. S11).

We also observed variations in electrophoretic mobility across the designs: PDN8 and PDN32 exhibit higher electrophoretic mobility, although their molecular weights (25.25 kDa and 25.68 kDa, respectively) are comparable to that of the natural protein A. One possible reason for the inconsistency between electrophoretic mobility and molecular weight might be local misfolding or conformational heterogeneity of the designs.

Together, the high expression levels and robust solubility observed across these designed proteins underscore the effectiveness of both the generating and selection approaches used by OCDesign.

##### (ii) IgG-binding affinity

We quantified the binding capacity of the designed proteins toward IgG antibody via ELISA. As shown in Fig. 3**c**, **d**, and Table S11, PDN4 and PDN8, the Pareto fronts selected in round 1, exhibit low binding affinities of 0.042 and 0.115, respectively. In contrast, the four designs in round 2 exhibit high binding affinity (PDN12: 1.40, PDN15: 2.19, PDN17: 1.84, PDN19: 0.331), with PDN15 even higher than that of the natural protein A. Three out of the five designs in round 3 also exhibit strong binding affinity (PDN26: 0.911, PDN31: 0.165, PDN32: 0.372). These results demonstrate that fixing key protein residues can increase IgG-binding affinity, and Pareto-front selection helps identify favorable trade-offs between protein solubility and IgG-binding affinity.

##### (iii) Structural consistency and stability

To ascertain whether the designed proteins adopt the intended structure, we characterized their secondary structure using circular dichroism spectroscopy. Here, we used PDN8 and PDN15 as representatives of the designs, which are Pareto fronts selected in rounds 1 and 2, respectively. As shown in Fig. 4**a, b**, PDN8 and PDN15 exhibit characteristic “W”-shaped spectra with minima at 208 nm and 222 nm. These spectral features reveal the dominant *α*-helical conformation, which is consistent with the triple-helix bundle architecture of protein A’s domains. An in-depth quantitative analysis of secondary-structure composition revealed that deviations between the proteins and the natural protein A were below 5% (Table S12).

We further performed differential scanning calorimetry (DSC) to compare the thermal stability of PDN15 with that of the control. As shown in Fig. 4**c**, PDN15 exhibits a thermal denaturation temperature (*T_m_*) of 57.2 °C, lower than that of the natural protein A (92.8 °C). These results suggest that while the current design strategy successfully preserves the structural topology, further refinement of hydrophobic core packing or inter-helical interactions may be required to enhance thermal resilience.

##### (iv) Alkaline resistance

To assess the alkaline resistance of the designed proteins, we exposed them in 0.5 mol/L NaOH and monitored their structural integrity using SDS-PAGE. Here, we focused on the designs with significant binding affinity, including PDN8, 12, 15, 17, 19, 26, 31, and 32.

As shown in Fig. 6 and Fig. S7, PDN8, PDN12, PDN17, PDN19, and PDN26 showed early signs of band diffusion after 12 hours and underwent complete degradation within 24 hours. PDN15 demonstrated slightly improved performance, with minor degradation at 24 hours and complete loss of integrity by 36–48 hours. In contrast, PDN31 and PDN32, the final designs from round 3, exhibited alkaline stability comparable to the control even after 48 hours. We also observed that PDN18 degraded after 36 hours, indicating its relatively low alkaline resistance compared to PDN31 and PDN32.

Taken together, these findings suggest that the designs acquired in round 3, PDN31 and PDN32, have high solubility and alkaline resistance, as well as appropriate IgG-binding affinity, making them promising substitutes for the natural protein A as fillers for antibody purification. Thus, the objective curriculum principle can effectively guide the design of multi-property proteins.

#### 1.2 Full-length proteins with inserted linkers exhibit enhanced flexibility and binding sites

We constructed full-length proteins from the designed domains in two ways: (i) by directly cascading four domains, and (ii) by cascading four domains with a linker inserted between adjacent domains. Here, each linker is a 5-mer peptide, “KVDAK”, derived from the natural protein A. To investigate the effects of inserted linkers, we used the designed domain PDN31 as an example to construct two full-length proteins, one with inserted linkers and the other without linkers, built protein-antibody complexes through docking only one antibody onto a certain domain, and then performed MD simulation analysis (200 ns) by running GROMACS [42–46] on these complexes.

From the 200,000 frames generated by GROMACS, we extracted 2,000 and clustered them using the GROMOS algorithm [42–46] with a 1.0 nm RMSD cutoff. As shown in Fig. S6**a, b**, the full-length protein with inserted linkers exhibited higher structural flexibility: it yielded five clusters, whereas the protein without inserted linkers yielded only four clusters.

We further investigated how many domains can capture IgG antibodies after immobilizing the full-length proteins on an ELISA plate. The underlying rationale is that when immobilizing the full-length protein on an ELISA plate, the plate might clash with antibodies docked to certain domains, thereby precluding their binding. Here, we assume the plate is immobilized at the site opposite the predicted IgG-binding interface of a selected domain. As shown in Fig. S6**c, d**, with inserted linkers, the full-length protein has two domains that can bind IgG antibodies, whereas the full-length protein without linkers has only one such domain. This finding suggests that inserting linkers might increase the likelihood of binding to IgG antibodies.

**Table S1:**
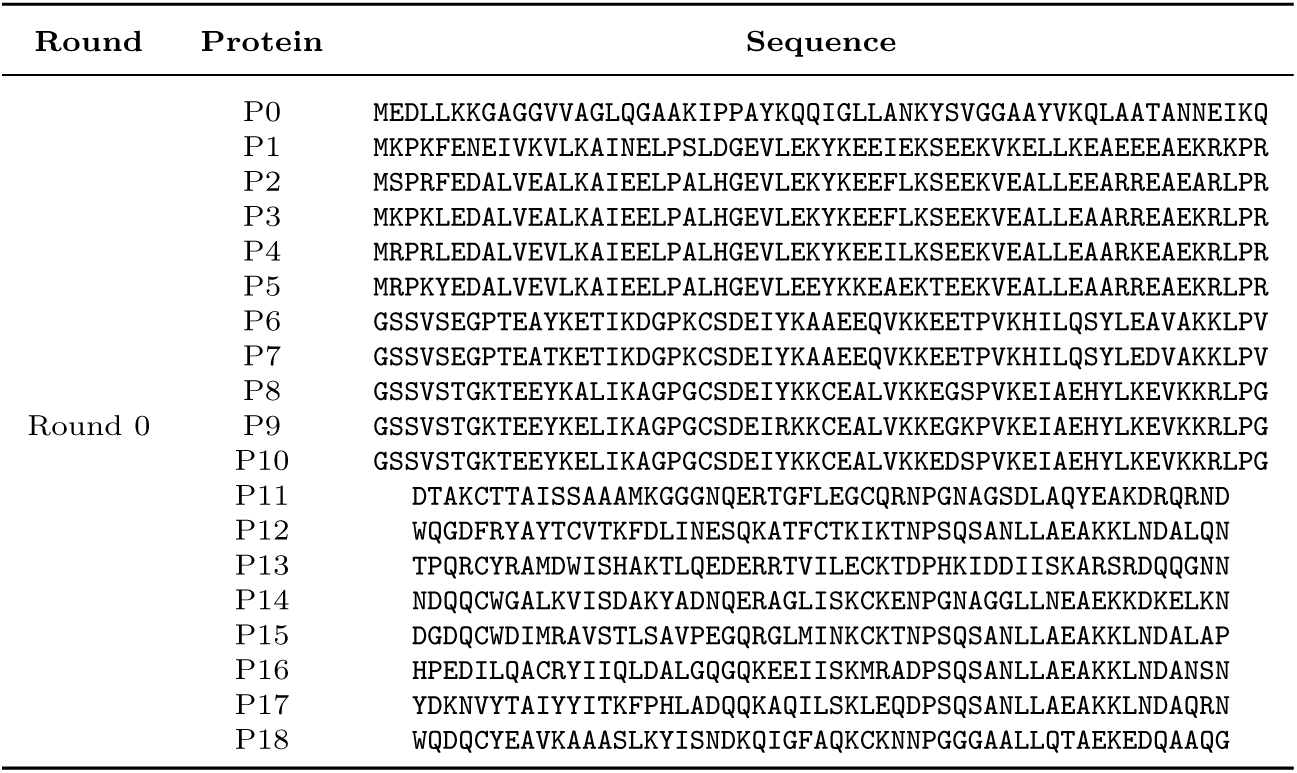
Designed proteins from the preliminary design campaign. Here, P0-10 were designed according to domain Z (PDB ID: 2SPZ; 58 residues) through optimizing three properties (i.e., solubility, structural self-consistency, and binding affinity), and P11-18 were designed according to domain B (PDB ID: 5U4Y; 53 residues) through optimizing only two objectives: solubility and structural self-consistency.

**Table S2:** *In silico* assessment of properties of the designed proteins from the preliminary design campaign. Here, TM-score describes the structural self-consistency, i.e., the similarity between the predicted structure of the designed proteins and the native structure of the natural protein A (B domain).

| Protein | Identity | TM-score |
| --- | --- | --- |
| P0 | 0.04 | 0.71 |
| P1 | 0.22 | 0.88 |
| P6 | 0.12 | 0.85 |
| P8 | 0.12 | 0.84 |
| P11 | 0.11 | 0.79 |
| P12 | 0.43 | 0.79 |

**Table S3:**
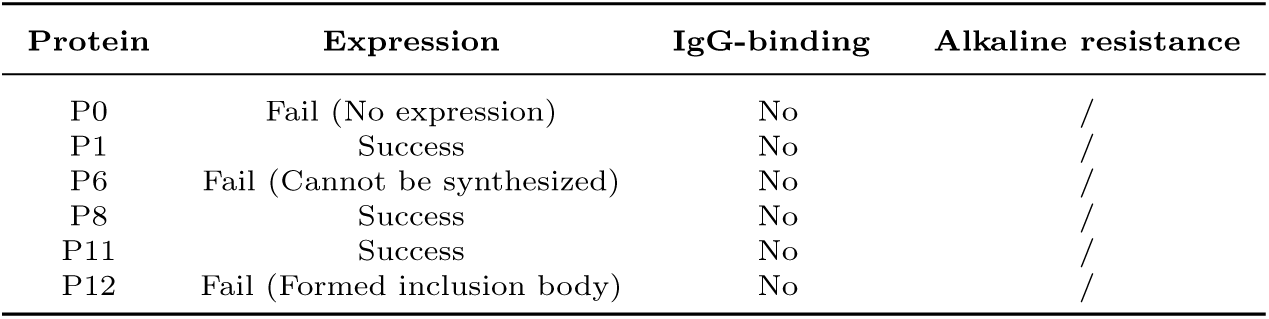
Wet-lab validation of the designed proteins from the preliminary design campaign.

**Table S4:**
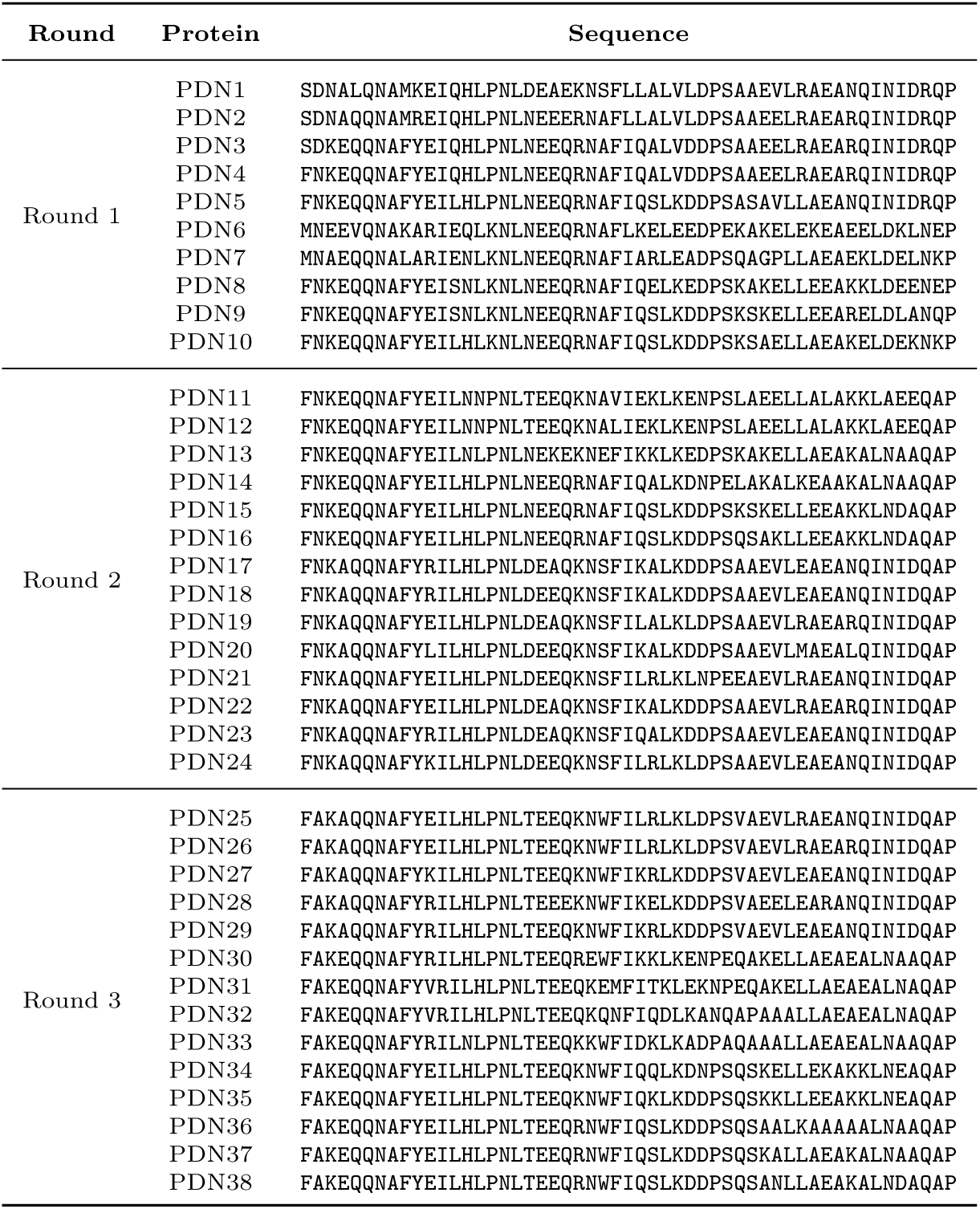
Designed proteins from the three rounds of OCDesign.

**Table S5:** *In silico* assessment of properties for designs from round 1. The Pareto fronts of the designs by NeuralMRF include PDN3, 4, and 5, while the Pareto fronts of the designs by ProDESIGN-LE include PDN6, 8, and 10. The visualization of these designs is provided in Fig. 2.

| Protein | Design method | Identity | TM-score | RMRD | Solubility |
| --- | --- | --- | --- | --- | --- |
| PDN1 | NeuralMRF | 0.43 | 0.943 | 1.01 | 0.500 |
| PDN2 | NeuralMRF | 0.53 | 0.963 | 0.995 | 0.563 |
| PDN3 | NeuralMRF | 0.68 | 0.965 | 1.00 | 0.902 |
| PDN4 | NeuralMRF | 0.72 | 0.963 | 0.660 | 0.920 |
| PDN5 | NeuralMRF | 0.83 | 0.959 | 0.676 | 0.934 |
| PDN6 | ProDESIGN-LE | 0.47 | 0.972 | 0.302 | 0.738 |
| PDN7 | ProDESIGN-LE | 0.60 | 0.966 | 0.316 | 0.654 |
| PDN8 | ProDESIGN-LE | 0.72 | 0.968 | 0.318 | 0.940 |
| PDN9 | ProDESIGN-LE | 0.75 | 0.963 | 0.330 | 0.926 |
| PDN10 | ProDESIGN-LE | 0.83 | 0.963 | 0.328 | 0.945 |

**Table S6:** Wet-lab validation of four selected designs from round 1.

| Protein | Expression | IgG-binding | Alkaline resistance |
| --- | --- | --- | --- |
| PDN3 | Success | No | No |
| PDN4 | Success | No | No |
| PDN6 | Success | No | No |
| PDN8 | Success | No | No |

**Table S7:** *In silico* assessment of properties for designs from round 2.

| Protein | Design method | Identity | TM-score | MM/GBSA (kcal/mol) | RMSD variance | Solubility |
| --- | --- | --- | --- | --- | --- | --- |
| PDN11 | ProDESIGN-LE | 0.679 | 0.930 | -65.7 | 0.098 | 0.919 |
| PDN12 | ProDESIGN-LE | 0.679 | 0.939 | -51.2 | 0.241 | 0.915 |
| PDN13 | ProDESIGN-LE | 0.736 | 0.952 | -83.3 | 0.074 | 0.919 |
| PDN14 | ProDESIGN-LE | 0.774 | 0.963 | -75.4 | 0.731 | 0.963 |
| PDN15 | ProDESIGN-LE | 0.925 | 0.959 | -66.0 | 0.100 | 0.972 |
| PDN16 | ProDESIGN-LE | 0.962 | 0.960 | -69.9 | 0.167 | 0.932 |
| PDN17 | NeuralMRF | 0.660 | 0.954 | -52.4 | 0.190 | 0.938 |
| PDN18 | NeuralMRF | 0.679 | 0.954 | -98.5 | 0.072 | 0.921 |
| PDN19 | NeuralMRF | 0.660 | 0.956 | -41.0 | 0.380 | 0.871 |
| PDN20 | NeuralMRF | 0.679 | 0.954 | -75.5 | 0.063 | 0.926 |
| PDN21 | NeuralMRF | 0.642 | 0.959 | -59.3 | 0.219 | 0.945 |
| PDN22 | NeuralMRF | 0.679 | 0.967 | -89.1 | 0.179 | 0.933 |
| PDN23 | NeuralMRF | 0.679 | 0.953 | -68.3 | 0.062 | 0.913 |
| PDN24 | NeuralMRF | 0.660 | 0.959 | -58.4 | 0.204 | 0.919 |

**Table S8:** Wet-lab validation of four selected designs from round 2.

| Protein | Expression | IgG-binding | Alkaline resistance |
| --- | --- | --- | --- |
| PDN12 | Success | Yes | No |
| PDN15 | Success | Strong | No |
| PDN17 | Success | Yes | No |
| PDN19 | Success | Yes | No |

**Table S9:** *In silico* assessment of properties for designs from round 3.

| Protein | Design method | Identity | TM-score | MM/GBSA (kcal/mol) | RMSD variance | Solubility |
| --- | --- | --- | --- | --- | --- | --- |
| PDN25 | NeuralMRF | 0.660 | 0.966 | -52.9 | 0.081 | 0.933 |
| PDN26 | NeuralMRF | 0.660 | 0.961 | -54.1 | 0.161 | 0.883 |
| PDN27 | NeuralMRF | 0.660 | 0.962 | -55.6 | 0.132 | 0.930 |
| PDN28 | NeuralMRF | 0.623 | 0.965 | -82.2 | 0.141 | 0.939 |
| PDN29 | NeuralMRF | 0.660 | 0.970 | -45.0 | 0.061 | 0.949 |
| PDN30 | ProDESIGN-LE | 0.698 | 0.967 | -55.3 | 0.110 | 0.941 |
| PDN31 | ProDESIGN-LE | 0.660 | 0.955 | -77.4 | 0.168 | 0.918 |
| PDN32 | ProDESIGN-LE | 0.717 | 0.968 | -56.5 | 0.076 | 0.927 |
| PDN33 | ProDESIGN-LE | 0.698 | 0.958 | -63.9 | 0.482 | 0.881 |
| PDN34 | ProDESIGN-LE | 0.792 | 0.953 | -73.1 | 0.570 | 0.970 |
| PDN35 | ProDESIGN-LE | 0.830 | 0.954 | -61.2 | 10.6 | 0.973 |
| PDN36 | ProDESIGN-LE | 0.830 | 0.967 | -62.9 | 0.174 | 0.924 |
| PDN37 | ProDESIGN-LE | 0.868 | 0.966 | -57.9 | 0.127 | 0.940 |
| PDN38 | ProDESIGN-LE | 0.925 | 0.965 | -58.4 | 0.097 | 0.965 |

**Table S10:** Wet-lab validation of five selected designs from round 3.

| Protein | Expression | IgG-binding | Alkaline resistance |
| --- | --- | --- | --- |
| PDN25 | Success | No | No |
| PDN26 | Success | Yes | No |
| PDN29 | Success | No | No |
| PDN31 | Success | Yes | Yes |
| PDN32 | Success | Yes | Yes |

**Table S11:**
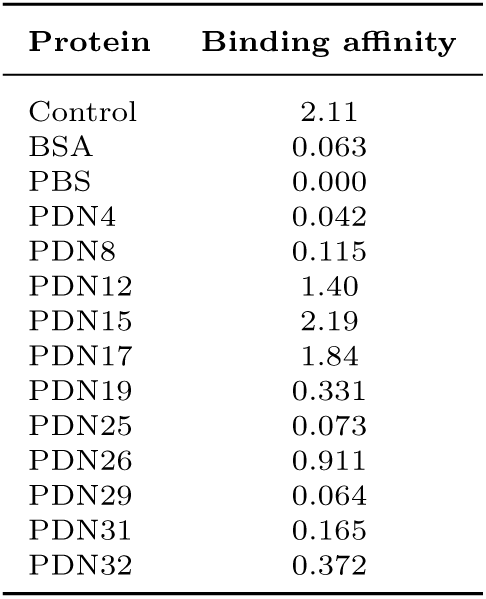
Characterizing binding affinity (OD_450_) of protein-IgG complexes using ELISA technology. Here, each complex consists of a designed protein and an IgG antibody.

**Table S12:** Structural characterization of PDN8, PDN15, and the natural protein A (Control) using circular dichroism (CD) technology. Here, NRMSD denotes the normalized root mean deviation between an experimental CD spectrum and its theoretical counterpart. The acquired CD spectra were analyzed using Spectra Manager software (JASCO co.).

| Protein | $\alpha$ helix | $\beta$ sheet | $\beta$ turn | coil | NRMSD |
| --- | --- | --- | --- | --- | --- |
| Control | 82.60% | 8.50% | 4.90% | 4.00% | 0.056 |
| PDN8 | 84.30% | 9.00% | 3.30% | 3.40% | 0.062 |
| PDN15 | 85.30% | 6.20% | 3.50% | 5.00% | 0.074 |

**Fig. S1:**
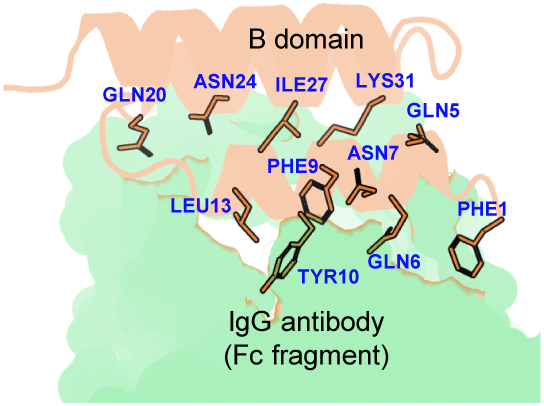
The binding of protein A to antibody IgG acquired from protein complex 5U4Y. Here, only domain B of protein A is shown for the sake of clarity. We also depict the key residues of protein A involved in its binding to IgG.

**Fig. S2:**
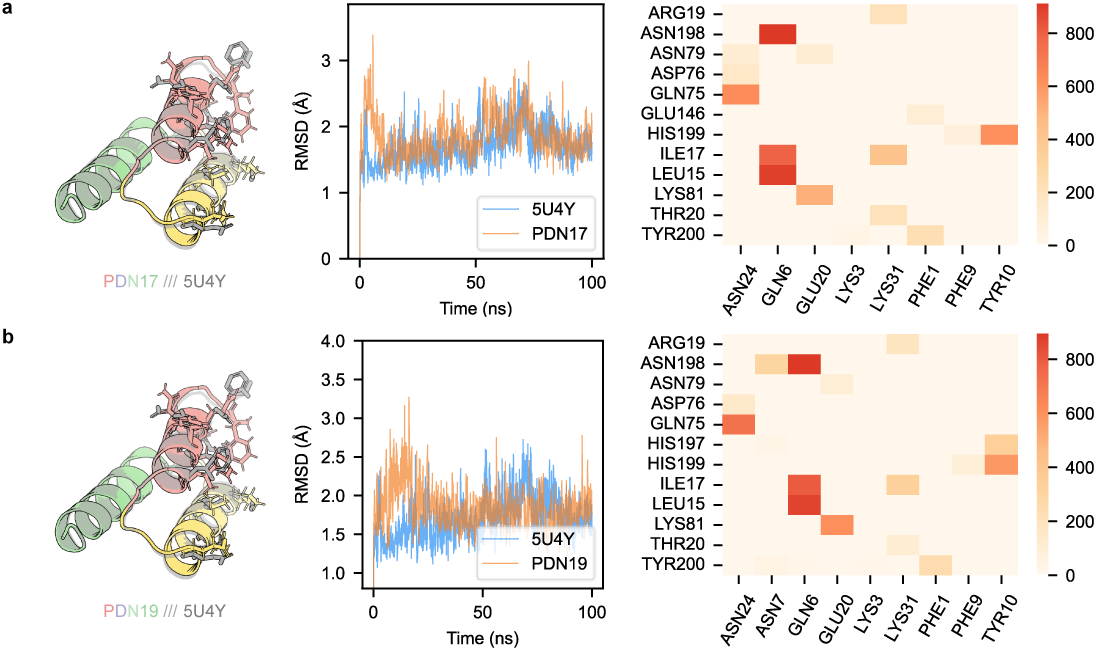
Molecular dynamics simulation analysis of the protein-IgG complexes constructed from PDN17 and PDN19. **a, b** MD simulation trajectories and key binding residues of protein-IgG complexes formed by PDN17 and PDN19. Here, the key binding residues are displayed as a matrix: columns correspond to IgG residues, and rows to design residues.

**Fig. S3:**
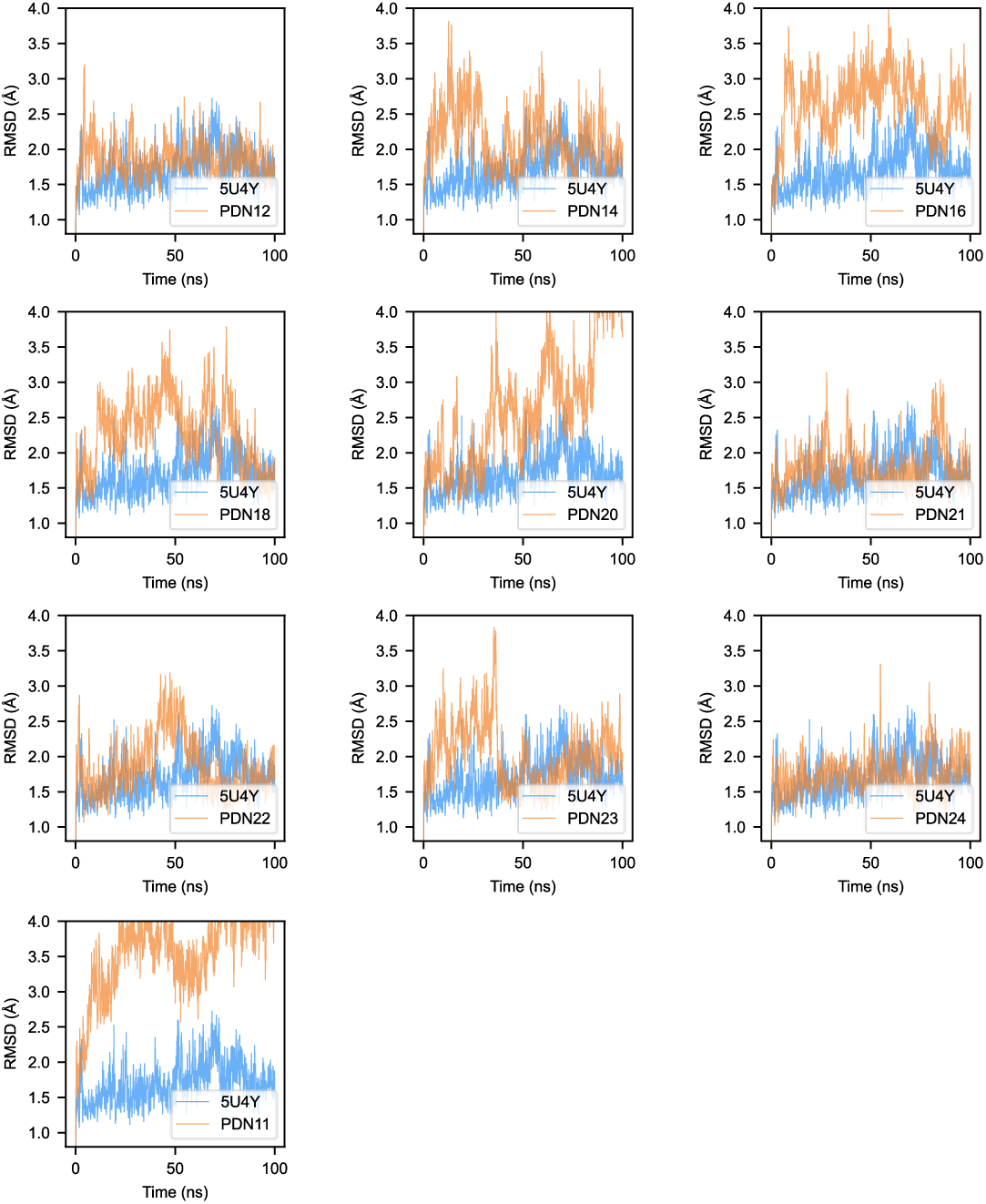
Molecular dynamics (100 ns) trajectory of the protein-IgG complexes, each complex consisting of a design from round 2. The designs selected for wet-lab validation, PDN17 and PDN19, are shown in Fig. S2.

**Fig. S4:**
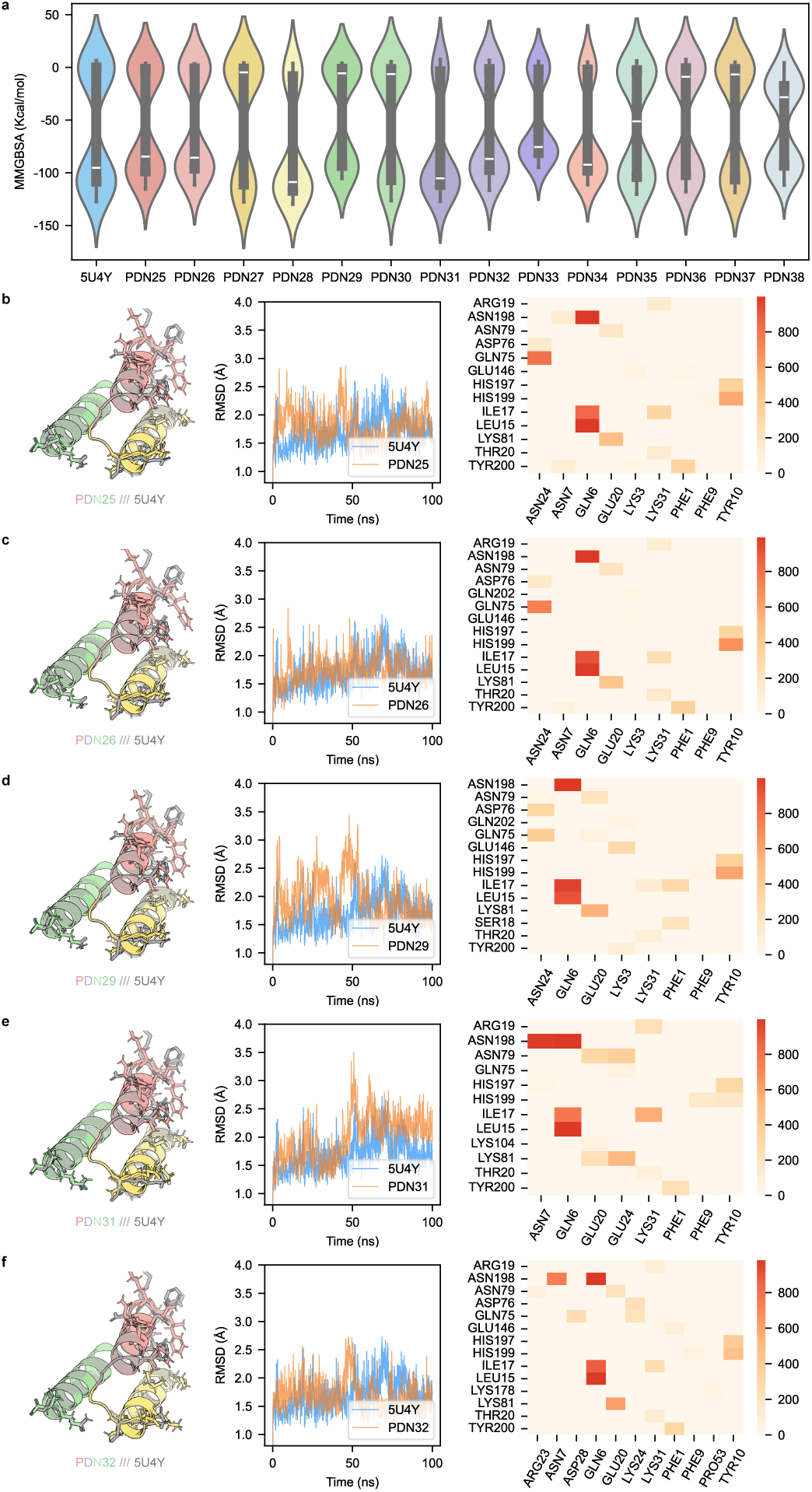
Molecular dynamics simulation analysis of protein-IgG complexes, each complex consisting of a round 3 design, including PDN26, PDN29, PDN31, and PDN32. **a,** Distribution of binding free energy (MM/GBSA) calculated from simulation results over 90–100 ns. **b-f,** MD simulation trajectories and key binding residues for complexes formed with native protein A, PDN25, PDN26, PDN29, PDN31, and PDN32. Key binding residues are shown as matrices: columns correspond to IgG residues, rows to residues of native protein A or the designs.

**Fig. S5:**
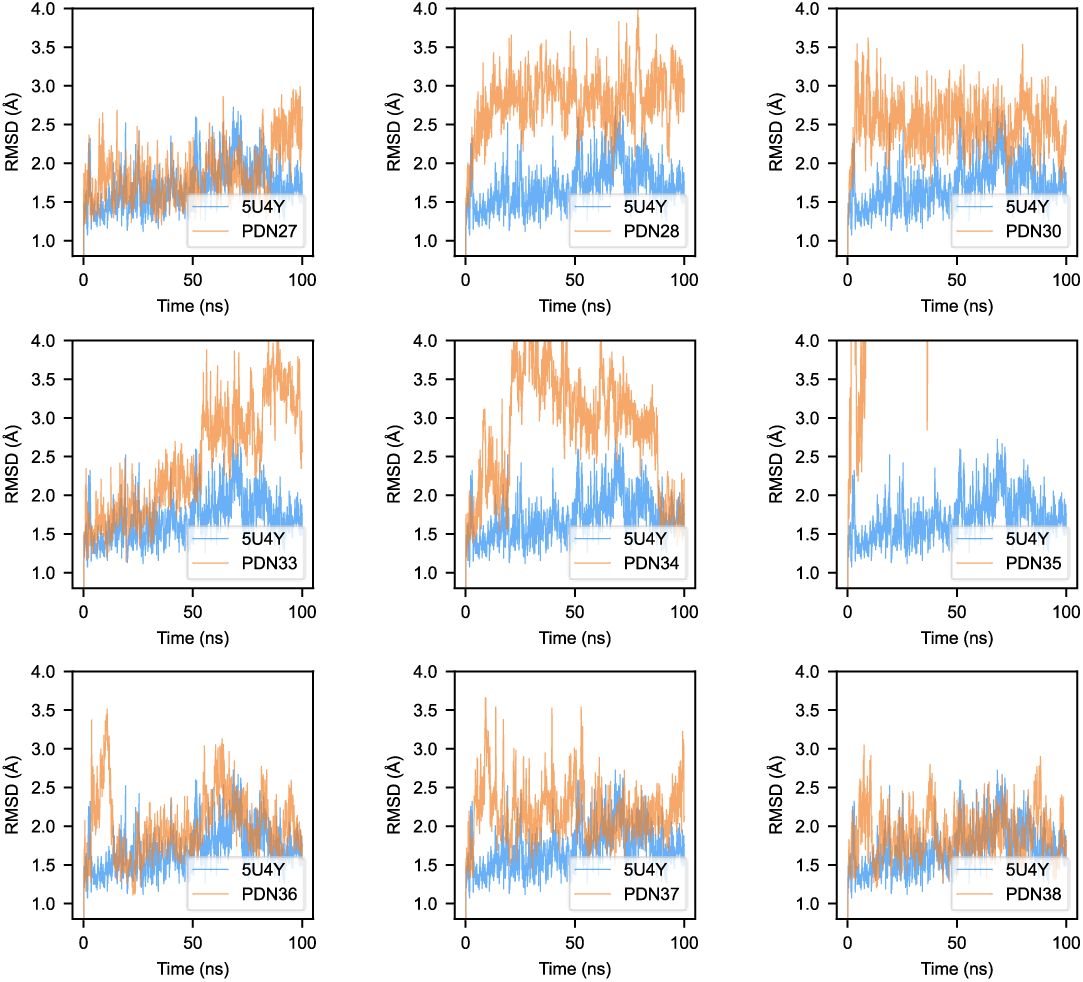
Molecular dynamics (100 ns) trajectory of the protein-IgG complexes, each complex consisting of a design from round 3. The designs selected for wet-lab validation, PDN26, PDN29, PDN31, and PDN32, are shown in Fig. S4.

**Fig. S6:**
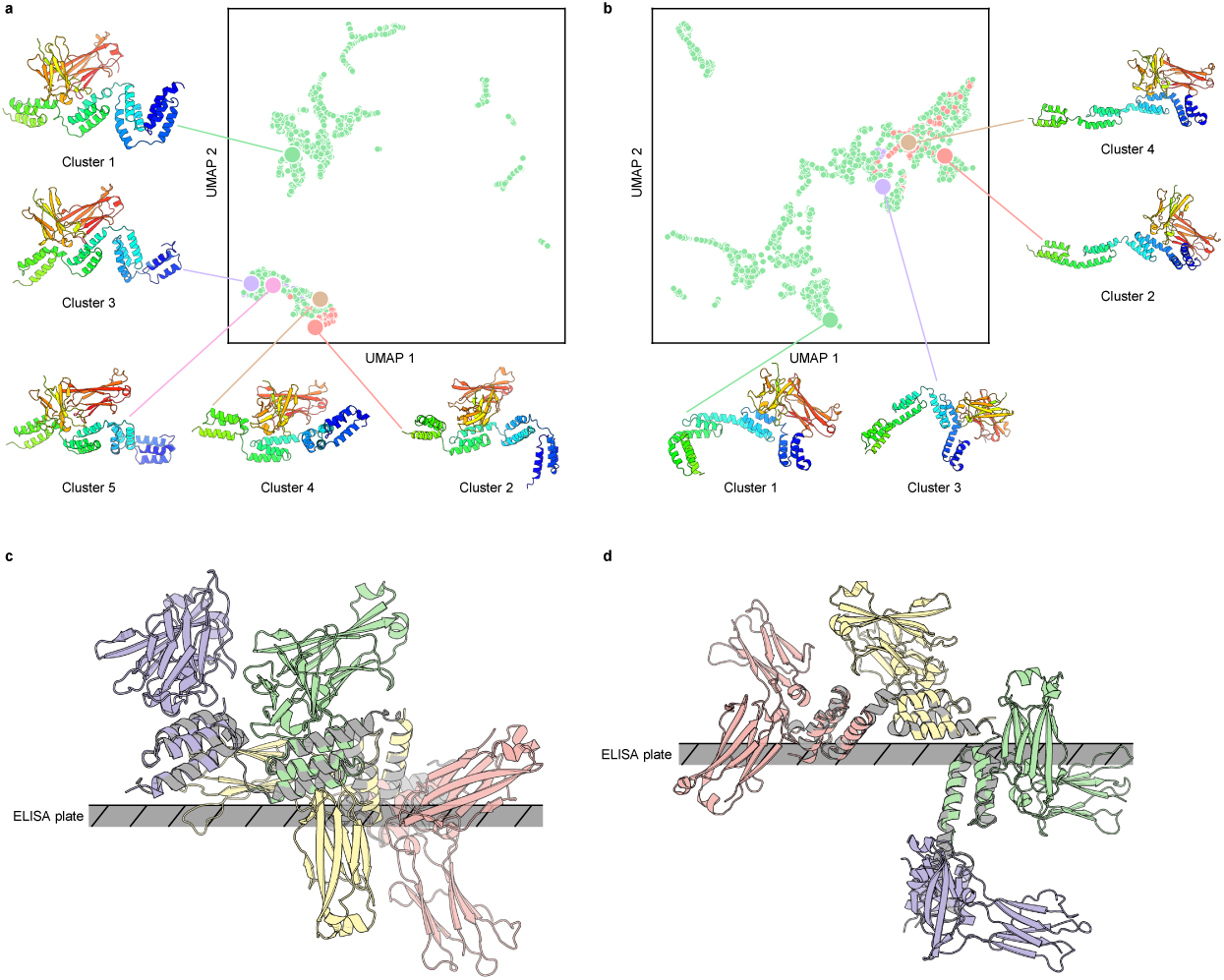
Investigating the effects of inserted linkers for constructing full-length proteins. We constructed full-length proteins from the designed domains in two ways: (i) by directly cascading four domains and (ii) by cascading four domains with a linker inserted between adjacent domains. Here, we use the design PDN31 as an example. The linker is a 5-mer peptide, “KVDAK”, excerpted from the natural protein A. **a, b,** Cluster-center structures from 200 ns GROMACS molecular dynamics simulations of predicted full-length PDN31-IgG complex with (a) or without (b) linkers, sampled over 2,000 frames. The linker-containing construct yielded five clusters, whereas the linker-free construct yielded four. **c, d,** Binding of IgG to the full-length protein with (c) or without (d) linkers when the full-length protein is immobilized on an ELISA plate (gray stripe). The linker-containing construct accommodates two IgG molecules, whereas the linker-free construct accommodates only one IgG.

**Fig. S7:**
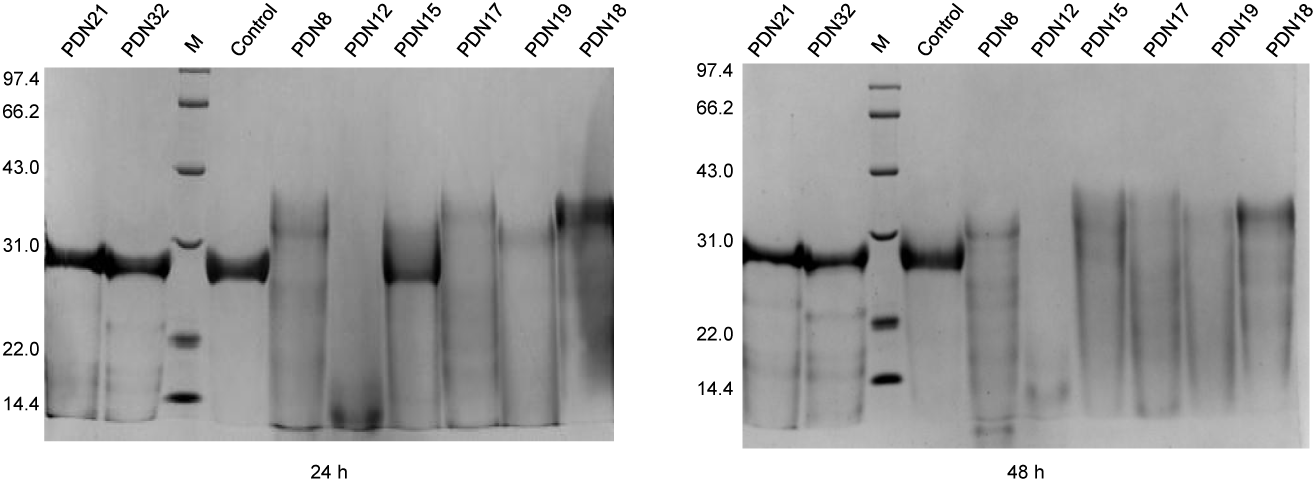
Characterizing alkaline resistance of the designed proteins using SDS-PAGE (treated with 0.5 mol/L NaOH after 24 and 48 hours). To investigate the alkaline resistance of the designed proteins, we continuously treated them with 0.5 mol/L NaOH solution (after 0, 12, 24, 36, and 48 hours) and examined their integrity using SDS-PAGE. The data acquired after 0-, 12-, and 36-hour treatments are provided in Fig.6.

**Fig. S8:**
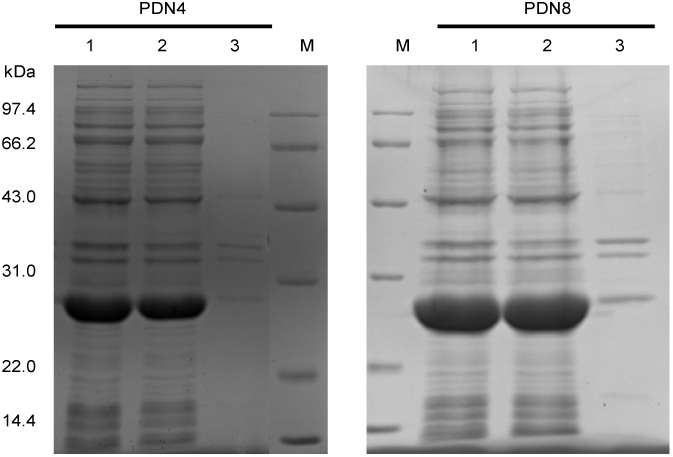
Characterizing expression and solubility of two round 1 designs, PDN4, and PDN8, using SDS-PAGE. The lanes are as follows. Lane M: Protein molecular weight marker (kDa). Lane 1: Whole cell lysate after induction. Lane 2: Soluble supernatant after cell disruption and centrifugation. Lane 3: Insoluble pellet after cell disruption and centrifugation.

**Fig. S9:**
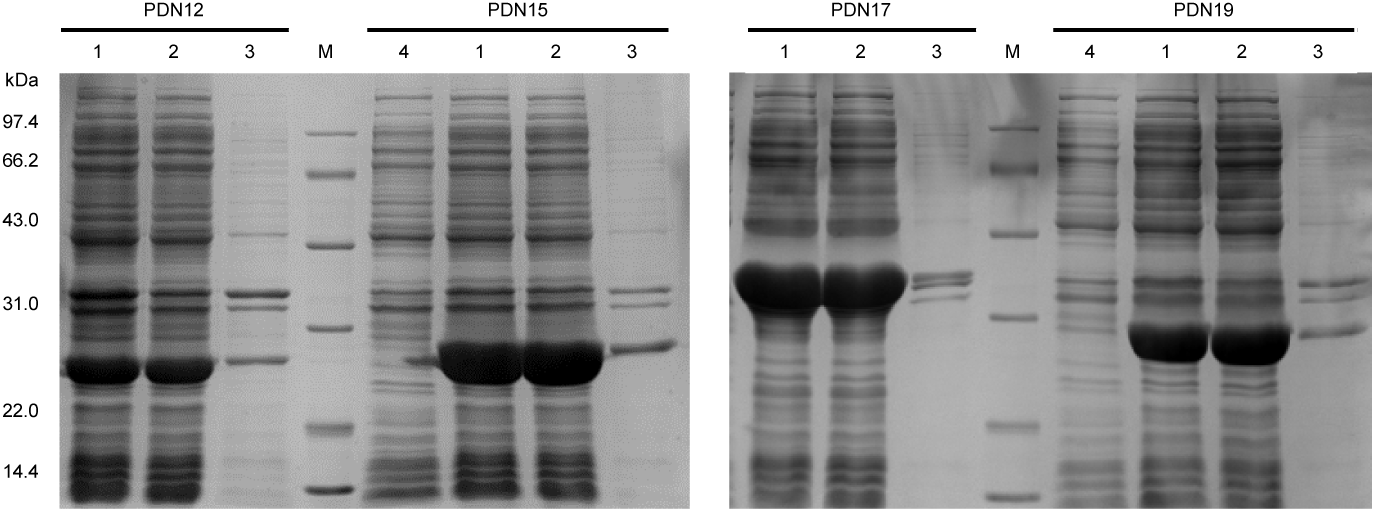
Characterizing expression and solubility of four round 2 designs, PDN12, PDN15, PDN17, and PDN19, using SDS-PAGE. The lanes are as follows. Lane M: Protein molecular weight marker (kDa). Lane 1: Whole cell lysate after induction. Lane 2: Soluble supernatant after cell disruption and centrifugation. Lane 3: Insoluble pellet after cell disruption and centrifugation. Lane 4: Whole cell before induction (negative control).

**Fig. S10:**
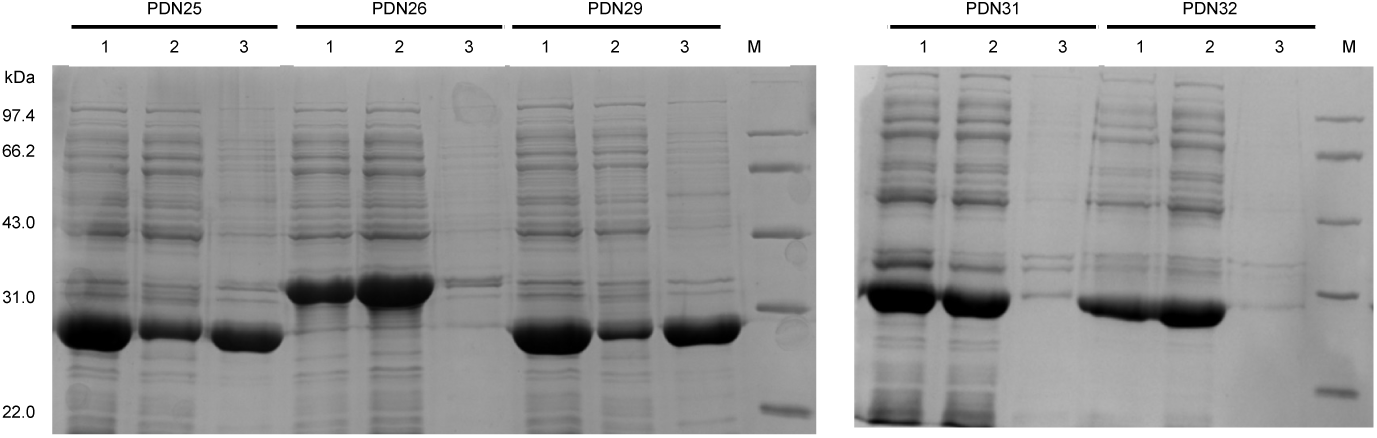
Characterizing expression and solubility of five round 3 designs, PDN25, PDN26, PDN29, PDN31, and PDN32, using SDS-PAGE. The lanes are as follows. Lane M: Protein molecular weight marker (kDa). Lane 1: Whole cell lysate after induction. Lane 2: Soluble supernatant after cell disruption and centrifugation.Lane 3: Insoluble pellet after cell disruption and centrifugation.

**Fig. S11:**
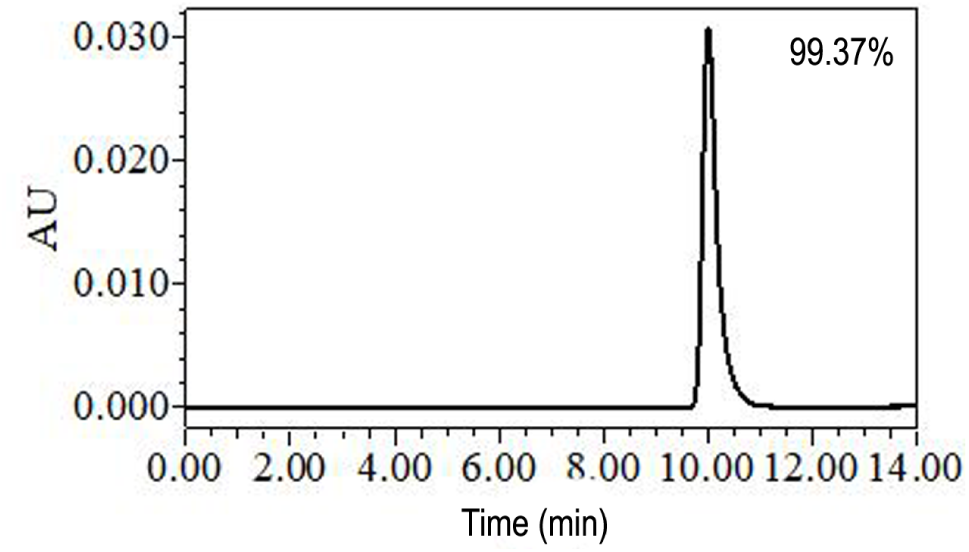
Characterizing purity of the design protein (PDN15) using SEC-HPLC. Size-exclusion high-performance liquid chromatography (SEC-HPLC) was used to assess the homogeneity and purity of the acquired protein PDN15. The chromatogram shows a single symmetric peak at a retention time of μ10 min, accounting for 99.37% of the total peak area, i.e., the purity of the acquired proteins reaches 99.37%.

### 2 Methods

#### 2.1 One-shot optimization for multi-property protein design

In the preliminary design campaign, we tested whether multiple desired properties could be achieved simultaneously in a single shot. We therefore implemented a one-shot optimization strategy, termed ProDESIGN-MCMC, in which solubility, structural self-consistency, and IgG-binding affinity were considered simultaneously in a single Markov chain Monte Carlo (MCMC) search.

Specifically, solubility was optimized by minimizing the negative pseudo-likelihood of the designed protein sequence, while structural self-consistency and IgG-binding affinity were jointly optimized by minimizing the cross-entropy between the distograms of the native complex and the predicted complex of the designed sequences with IgG. This strategy differs from OCDesign, where these objectives are introduced sequentially across design rounds.

ProDESIGN-MCMC combines two components: the protein language model ProGPT (trained by us and unpublished) to evaluate sequence naturalness and solubility, and the protein structure prediction model AlphaFold2 to enforce structural self-consistency and IgG-binding-compatible geometry of the designed sequences.

##### One-shot sequence design

We design a protein sequence for a target backbone as follows:

1. Initialization: Two methods are available to obtain a starting sequence of the same length as the target backbone. The first method generates multiple sequences and selects the best as the starting sequence. The second method randomly generates a single sequence. Both are valid because this MCMC has a stationary distribution; in theory, the result is independent of the starting sequence. In practice, for efficiency, we randomly select the starting sequence.
2. Iteration update: Each iteration consists of two steps. *i)* Sample update position: For the current designed sequence, we predict pLDDT using AlphaFold2, then sample an update position according to the pLDDT of each residue. *ii)* Sample residue: For the selected position, we compute the probabilities for the 20 candidate residues according to Equation S2.
3. Stop condition: We stop a design trial based on the number of iterations. In practice, we balance design time and performance by performing 4 *× L* iterative updates (*L* is the sequence length).

##### Objective function of joint optimization

ProDESIGN-MCMC is a method for designing protein sequences for a given backbone structure. We formulate the problem as maximizing the probability P(seq *|* str), and decompose it using Bayes’ rule as follows:

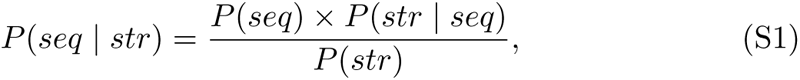

where *P*(*seq*) is the probability that the sequence follows natural protein sequence rules, evaluated by our trained protein language model ProGPT; *P*(*str | seq*) is the probability that the sequence folds into the target backbone structure, evaluated by the distogram loss between the designed sequence and the target backbone; and *P*(*str*) is a constant representing the probability of the input backbone structure in the space of native backbone structures.

To ensure the computability of *P*(*seq|str*), we using the logarithm to calculate, and update the optimization function as follows:

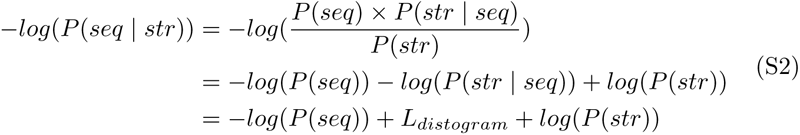

In the preliminary design campaign (round 0), we used two target backbone structures, including the Z domain (2SPZ) and the B domain-IgG Fc complex (5U4Y). We changed the design target to ensure the IgG-binding affinity of the designs by constraining the binding position between the designs and the IgG Fc in the native format.

#### 2.2 MRF-based protein design method NeuralMRF

The input to NeuralMRF is the three-dimensional (3D) structure of a protein, and the output is the corresponding amino acid sequence. Its architecture consists of two primary components: an encoder based on a message-passing neural network (MPNN) and an MRF-based decoder.

##### 2.2.1 Basic idea of NeuralMRF

Built upon the ProteinMPNN framework, our encoder is designed to extract spatial features from the protein backbone (Details in Algorithm 1). We construct the graph neural network (GNN) graph using 48 nearest neighbors per residue based on *Cα*-*Cα* distances. Edge features are parameterized by 16 Gaussian Radial Basis Functions (RBFs) spanning 2Å to 22Å based on interresidue distances between heavy atoms (*Cα, Cβ, N, O*), while node features are zero-initialized.

For the decoding component, we employ an MRF-based decoder. Specifically, the MRF decoder is implemented using two separate Multi-Layer Perceptrons (MLPs): one for decoding the single-site amino acid probabilities and the other for capturing the pairwise coupling probabilities between amino acid residues. Compared to auto-regressive decoders, this approach offers several advantages. During training, the model provides direct supervision over amino acid pair probabilities. During inference, the MRF-based decoder exhibits superior efficiency as it enables simultaneous updates across all positions in the protein sequence, rather than following a sequential residue-by-residue generation process.

**Fig. S12:**
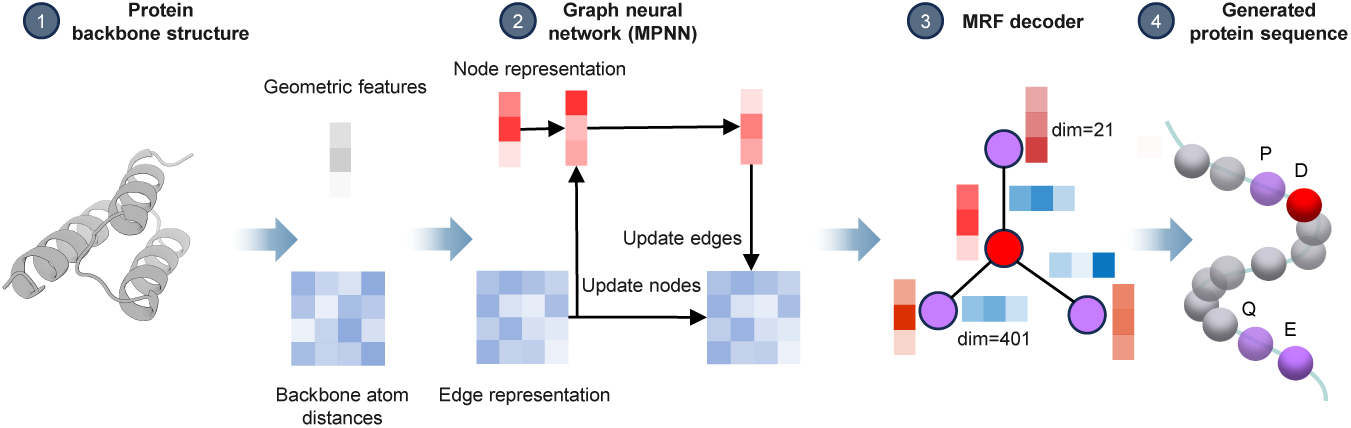
Overview of NeuralMRF. Given an input protein backbone structure, NeuralMRF designs a protein sequence that folds into the input backbone structure. First, it parses geometric features from the backbone to obtain node and pair representations for the GNN encoder module. Then, the GNN encoder updates the node and edge representations. Finally, the MRF decoder module decodes the protein sequence from the encoded representation.

NeuralMRF is a computational framework for protein sequence design grounded in the MRF principle. The implementation of the NeuralMRF is based on the ProteinMPNN to improve the performance in recovering the protein sequence and the explainability (Fig. S12). The basic idea of NeuralMRF is described below:

##### 2.2.2 Model architecture

NeuralMRF utilizes an encoder-decoder architecture. The encoder of the NeuralMRF is an MPNN that aims to capture the geometric features of the input protein backbone. The decoder of the NeuralMRF subsequently transforms these structural representations into an MRF formulation, from which the optimal protein sequence is derived.

##### 2.2.3 Encoder of NeuralMRF

The encoder module employs an MPNN comprising three layers. This module maps the input protein structure into two distinct latent spaces: node features (*V*) and pairwise edge features (*E*).

- Node Features (*V*): Initialized as all-zero vectors.
- Pairwise Features (*E*): Following the conventions of ProteinMPNN, distances between heavy atoms (*Cα, Cβ, N,* and *O*) for the 48 nearest neighbor residues are encoded using 16 RBFs uniformly spaced from 2Å to 22Å.
- Graph Sparsity: The GNN graph is constructed using the 48 nearest neighbors based on *Cα*–*Cα* distances. Edges exceeding a distance threshold of 15Å are masked to maintain structural locality.

##### 2.2.4 Decoder of NeuralMRF

The decoder projects the latent node and pair representations into the single-body and pair-body terms of an MRF. This explicit mapping facilitates greater interpretability, as the parameters correspond directly to the statistical potentials of the sequence-structure relationship. The module utilizes two linear layers to transform the representations:

- Node projection: Maps 128-dimensional representations to 21 dimensions (the 20 standard amino acids plus an uncertain token).
- Edge projection: Maps 128-dimensional representations to 441 dimensions (representing 21 *×* 21 possible residue-pair interactions).

###### Algorithm 1 MPNN encoder layer of NeuralMRF

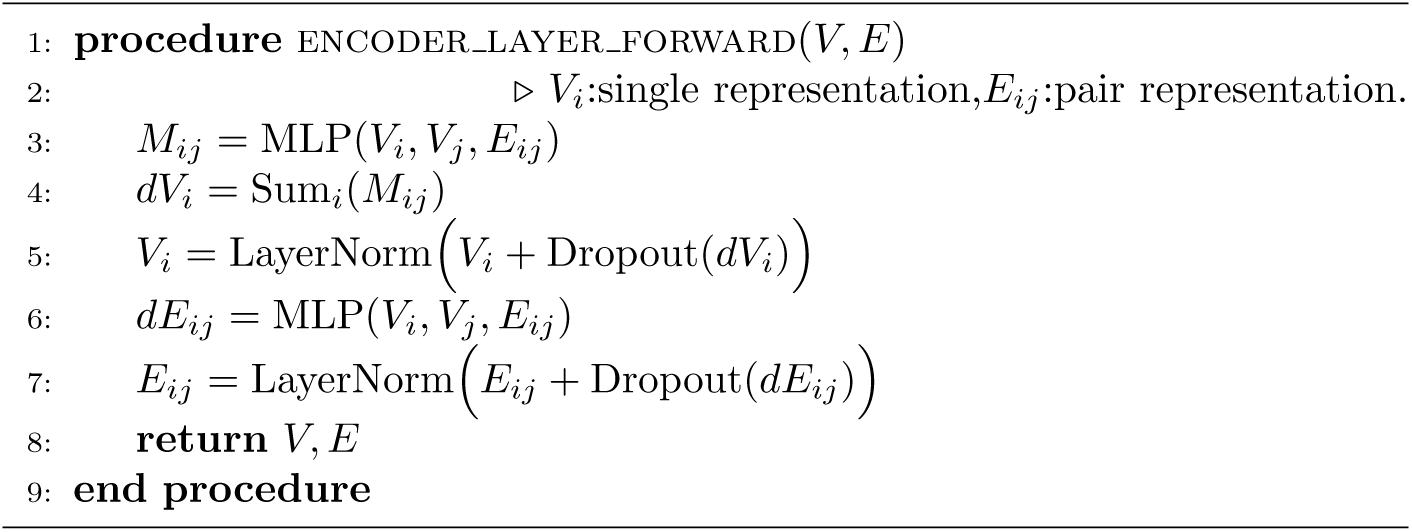

The definition of the MPNN encoder layer is as follows.

##### 2.2.5 Training data

NeuralMRF was trained and validated using 19,752 high-resolution single-chain structures obtained from the CATH 4.2 40% non-redundant dataset. The dataset was split the dataset into train, validation, and test sets (80/10/10).

##### 2.2.6 Test set

To evaluate the generalization and accuracy of the model, we assessed NeuralMRF on 92 protein structures from the 15th Critical Assessment of protein Structure Prediction, and the single-chain protein structures from the Continuous Automated Model Evaluation that were released between January 2022 and October 2022, filtered for a maximum length of 600 residues.

##### 2.2.7 Training process

We trained NeuralMRF using two objective functions: a cross-entropy loss for direct sequence recovery (*L*_seq_) and a maximum likelihood loss for the MRF (*L*_MRF_), which maximizes the joint probability of amino acid pairs. Additionally, an *L*_2_ regularization term was incorporated into the total loss to prevent overfitting (Equation S3). To address the computational intractability of training the MRF using full likelihood, we followed previous work and adopted a pseudo-likelihood approximation.

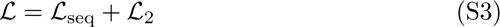

The NeuralMRF model was optimized using the Adam optimizer [47] with a learning rate of 0.004, *β*_1_ = 0.9, and *β*_2_ = 0.999. We conducted the training on four Nvidia RTX 3090Ti GPUs. To maintain a global batch size of 64, the local batch size was set to 4 per GPU with a gradient accumulation of 4 steps.

#### 2.3 *In silico* assessing multiple properties of design

In the dry-lab step of the design process, we assessed multiple properties of the designs, including solubility, structural self-consistency, stability, and binding affinity, to subsequently select Pareto-optimal designs that optimize these properties. These properties are assessed as follows:

##### Solubility

The solubility of designed proteins was assessed by running SoluProt, which returns a value within the range of [0, 1].

##### Structural self-consistency

We evaluated the structural self-consistency of the designs by computing the similarity (measured using TM-score) between their predicted structures and the desired structure. A higher TM-score indicates that the designed protein folds more accurately into the desired structure.

##### Stability

To assess the stability of a designed protein, we performed molecular dynamics simulations (100 ns) on the protein-antibody complexes: Each complex consists of a designed protein and the target antibody, and its tertiary structure was built through docking the AlphaFold2 prediction of the design with the target antibody [48] and was used as the initial structure for MDS.

We measured the stability of each constructed complex using the variance of the C*_α_* root-mean-square deviation (RMSD) between the complex’s initial structure and its variants along MD trajectories. As shown in Table S7, and S9, some complexes exhibit a considerable variance during MD simulation, while others change pretty little. For example, the complex comprising the design PDN35 deviates from its initial structure by up to 18.6Å and ultimately adopts a structure conformation with an RMSD of 11.7Å. In contrast, the complex comprising the design PDN26 deviates from its initial structure by less than 0.061 during the entire simulation, suggesting stronger binding stability of these complexes. We used the complex comprising the wild-type protein A and the target antibody (PDB: 5U4Y) as a control, which exhibits a low variance of 0.087.

##### Binding affinity

The binding affinity between a design and the target antibody was also assessed using MD simulation of the complex comprising these two proteins. Unlike stability calculations that consider all frames in an MD trajectory, binding affinity calculations focus only on the stable frames. Specifically, we extracted the final 100 frames (90-100 ns) from the MD simulation, during which the system exhibits considerable stability, and calculated the average binding free energy of these frames using the Prime MM/GBSA method. As shown in Table S7 and S9, the complex 5U4Y exhibits a low binding free energy of -57.46 kcal/mol. Some designs, e.g., PDN21, PDN24, and PDN32, achieve a binding free energy at the same level, indicating strong interactions between these designs and the target antibody.

#### 2.4 Protein expression and purification

Each expression plasmid was transformed into *Escherichia coli* BL21 (DE3) (CD601-02, TransGen Biotech). Protein expression was induced at an OD_600_ of 0.6–0.8 with 0.25 mmol/L IPTG for 4 h at 37 °C in LB medium containing 100 µg/mL kanamycin. Cells were harvested and sonicated in lysis buffer (20 mmol/L Na_3_PO_4_, 20 mmol/L imidazole, 500 mmol/L NaCl, pH 7.4). The soluble fraction was purified by Ni-affinity chromatography and eluted in a buffer containing 300 mmol/L imidazole, pH 7.4. After Ni-affinity purification, the sample was desalted, and the buffer was exchanged to 20 mmol/L Tris-HCl, pH 8.0, for subsequent anion exchange chromatography. The anion exchange column was equilibrated with 20 mmol/L Tris-HCl (pH 8.0). After washing with equilibration buffer, the target protein was eluted with 50 mmol/L Tris-HCl (pH 8.0) containing 500 mmol/L NaCl. Fractions were collected based on UV absorbance and analyzed by SDS-PAGE.

#### 2.5 ELISA analysis of protein–antibody binding

The binding affinity between the designed proteins and antibodies was evaluated by ELISA. Briefly, the newly designed protein molecules were diluted in carbonate buffer solution (CBS) and coated onto 96-well plates at 50 ng per well. The plates were incubated overnight at 4 °C. After washing three times with 100 µL PBST per well, the plates were blocked with 3% bovine serum albumin (BSA) at 37 °C for 1 h to reduce nonspecific binding.

After three additional washes with 100 µL PBST per well, the plates were incubated with horseradish peroxidase (HRP)-conjugated anti-rat IgG antibody at a dilution of 1:10,000 for 1 h at 37 °C. The plates were then washed thoroughly with PBST, followed by the addition of tetramethylbenzidine substrate solution (TMB; Solarbio). The colorimetric reaction was developed at 37 °C for 10 min in the dark and terminated with 50 µL sulfuric acid (2 mol/L) per well. The absorbance at 450 nm was measured using a microplate reader (Bio-Rad, Hercules, CA, USA).

Wells coated with BSA instead of the designed proteins were used as negative controls to evaluate nonspecific antibody binding.

#### 2.6 SDS-PAGE analysis to examine alkaline resistance

To evaluate the alkaline stability of the designed proteins, protein samples were prepared at a final concentration of 0.5 mg/mL in the presence of 0.5 mol/L NaOH. Control samples were prepared by adding an equal volume of ultrapure water instead of NaOH. The samples were incubated at room temperature, and aliquots were collected every 12 h. At each time point, 12% SDS-PAGE analysis was performed. All incubations were performed under sterile conditions.

#### 2.7 Circular dichroism spectroscopy for structural characterization

The circular dichroism spectra of the designed proteins and control protein (Control, US10501557B2, Date of Patent: Dec. 10, 2019) were recorded by a Japanese JASCO J-1500 Circular dichroism spectrometer using a 1 cm path length quartz cell. Protein solutions were added to 50 mmol/L PB, 5 mmol/L NaCl (pH 7.4) to give a final concentration of 5 mmol/L. Scans were recorded at 0.5 nm intervals from 190 to 260 nm at a rate of 60 nm/min, with a sensitivity of 100 millidegrees and a response time of 1 s. All spectra were recorded at 25 °C. The chromatographic signal from protein-free buffer was subtracted from each spectrum. Each spectrum was normalized to molar ellipticity (*θ*) using the mean weight residue and concentration. CD spectra were then converted to mean residue molar ellipticities.

#### 2.8 Analyzing protein thermal stability

Thermal stability of the proteins was assessed using the DSC TA Q2000 instrument. The temperature range was set from 30 °C to 120 °C. Protein samples were equilibrated at 30 °C in a nitrogen environment and then heated at a rate of 5 °C per minute to 120 °C without holding. Subsequently, the samples were cooled back to 30 °C at the same rate. Data was collected throughout the experiment, and thermal stability analysis graphs were generated using Origin software.

#### 2.9 Protein preparation and amino acid mutation

The co-crystal structure of Protein A-antibody (PDB ID: 5U4Y) was downloaded from the Protein Data Bank and prepared using Schrödinger suite (version 2021-1). For each mutant Protein A, amino acid residues at the designed sites were mutated according to the designed sequences (Supplementary Table S4), using NeuralMRF and ProDESIGN-LE. For example, in PDN9, residues at Q36, A38, N39, and A42 were mutated to K36, K38, E39, and E42, respectively. The resulting complex was PDN4-antibody complex. All protein-antibody complexes were preprocessed using the Protein Preparation Wizard [49] for complex optimization and minimization. Complex minimization was performed with the OPLS4 force field [50], with an RMSD for crystallographic heavy atoms at 0.3Å.

#### 2.10 Molecular dynamics simulations

Molecular dynamics simulations were conducted to evaluate the stability of the protein-antibody complexes. All systems were analyzed by the Desmond module [34] of Schrödinger suite for 100 ns simulation time. Preliminary tests showed that simulations of 200 ns yielded similar results to 100 ns; thus 100 ns was sufficient to capture significant conformational changes and save computational resources. System Builder module was used to prepare the complex systems for subsequent simulations. The atomic framework was solvated in an SPC [35] water model with orthorhombic boundary conditions for a 10Å buffer region. Counter ions were added to neutralize the system, and 0.15 M NaCl was added. The NPT ensemble (Isothermal-Isobaric ensemble) was applied to maintain a constant temperature [36] of 300 K and a pressure [37] of 1.01325 bar using the OPLS4 force field. A hybrid energy minimization algorithm with 1000 steps of steepest descent was applied, followed by conjugate gradient algorithms. Subsequently, the simulation trajectories were analyzed with the Simulation Event Analysis tool to compute the C*α* RMSD trajectory plots.

#### 2.11 Calculating MM/GBSA of the designed proteins

The Prime Molecular Mechanics/Generalized Born Surface Area (MM/GBSA) module in Schrödinger suite [51–53] was used to calculate the binding free energy of protein-antibody complexes from MD trajectory frames. In detail, MM/GBSA binding free energies were computed using the last 100 frames (90-100 ns) of the 100 ns MD simulation after the system reached equilibrium. The complex was refined in the OPLS4 force field adopting a VSGB solvation model [54]. The binding affinity of the protein-antibody complex was studied in terms of binding free energy (Δ*G*_bind_). Δ*G*_bind_ has mainly three influence factors: gas phase free energy (Δ*G*_MM_), solvation free energy (Δ*G*_GB_) and change in system entropy (Δ*G*_SA_). The relationship of the three factors is as follows: Δ*G*_bind_ = Δ*G*_MM_ + Δ*G*_GB_ + Δ*G*_SA_. The final values were obtained by averaging the results from the selected frames.

